# From vacuole to cytosol – Disruptive invasion triggers cytosolic release of *Salmonella* Paratyphi A and subsequent cytosolic motility favors evasion of xenophagy

**DOI:** 10.1101/2022.07.24.501230

**Authors:** Felix Scharte, Rico Franzkoch, Michael Hensel

## Abstract

*Salmonella enterica* is a common foodborne, facultative intracellular enteropathogen. Typhoidal *S*. *enterica* serovars like Paratyphi A (SPA) are human restricted and cause a severe systemic disease, while many *S*. *enterica* serovars like Typhimurium (STM) have broad host range, and in human hosts usually lead to self-limiting gastroenteritis. There are key differences between typhoidal and non-typhoidal *Salmonella* in pathogenesis, but underlying mechanisms remain largely unknown. Several genes encoding *Salmonella* pathogenicity island (SPI) effector proteins are absent or pseudogenes in SPA. Expression of virulence and metabolism genes show differential expression compared to STM. The intracellular transcriptomic architecture and phenotypes during presence in epithelial cells were recently described. Surprisingly, induction of motility, flagella and chemotaxis genes showed distinct expression patterns in intracellular SPA vs. STM and led to cytosolic motility of SPA. This study applies single cell microscopic analyses approaches to investigate the triggers and cellular consequences of cytosolic motility. Live cell imaging (LCI) revealed that SPA invades host cells in a highly cooperative manner. Extensive membrane ruffling at the invasion site leads to increased membrane damage in the nascent SCV with subsequent cytosolic release. After release into the cytosol, motile bacteria showed same velocity as under culture conditions used for infection. Reduced capture of SPA by autophagosomal membranes was observed by LCI and electron microscopy. Our results reveal flagella-mediated cytosolic motility as possible xenophagy evasion mechanism that could drive disease progression and contributes to dissemination of invasion-primed SPA during systemic infection.

**Importance:** Intracellular pathogens are commonly adapted to life in host cells either in a pathogen-containing vacuole, or free in host cell cytosol. However, transitions between these lifestyles are possible and demand specific adaptations, especially to avoid recognition and killing by host cell-autonomous immune defense. *Salmonella enterica* serovar Paratyphi A (SPA) belongs to typhoidal *Salmonella* able to cause live-threatening systemic infections in human hosts. We observed that SPA invades host cells in a way that often results in damage of the nascent vacuole and release of SPA in host cell cytosol. Here, SPA deploy flagella-mediated motility for rapid locomotion within infected cells. We demonstrate on single cell level that flagella-mediated motility enables evasion of xenophagic capture and control by the host cells. SPA uses a novel form of intracellular motility to successfully colonize human host cells.

## Introduction

Life inside mammalian host cells is a common virulence strategy of microbial pathogens and various forms of adaptation to life within host cell cytosol or pathogen-containing vacuoles (PCV), as well as exit strategies from host cells occur.

*Salmonella enterica* is a ubiquitous, invasive and facultative intracellular pathogen. Oral uptake of *S. enterica* by contaminated food or water causes infectious diseases ranging from self-limiting gastroenteritis to systemic infections with often fatal outcome (Dougan & Baker, 2014). While non-typhoidal *Salmonella* (NTS) serovars, such as *S. enterica* serovar Typhimurium (STM) often have broad host range, typhoidal *Salmonella* (TS) serovars such as *S*. Typhi (STY) or *S*. Paratyphi A (SPA) are characterized by adaptation to primate hosts in which typhoid or paratyphoid fever are important systemic diseases. The strict host adaptation limits studies of virulence mechanism of TS, and frequently infection models of STM in susceptible mouse strains are used as surrogate to investigate systemic *Salmonella* infections. However, SPA, STY and other TS are distinct from NTS by presence of increased accumulation of pseudogenes (Holt et al., 2009; McClelland et al., 2004), additional virulence factors as Vi capsule (STY) or typhoid toxin (STY, SPA), and distinct regulation of expression of virulence functions (Cohen et al., 2022; Reuter et al., 2021).

Both NTS and TS invade non-phagocytic mammalian cells, such as epithelial cells, by trigger invasion mediated by translocation of effector proteins by the *Salmonella* pathogenicity island 1 (SPI1)-encoded type III secretion system (T3SS). Host cell invasion is considered to initiate the intracellular lifestyle of *Salmonella*, and allows to breach epithelial barriers. Translocation of SPI1-T3SS effector proteins also evokes strong proinflammatory responses of epithelial cells, leading to intestinal inflammation, a hallmark of gastroenteritis by NTS. While SPI1-T3SS also mediates invasion by SPA and STY, intestinal inflammation usually is absent and other routes of entry appear to be used to breach epithelial barriers of the intestines in order to reach systemic sites (reviewed in Dougan & Baker, 2014).

Compiling data on the intracellular lifestyle in mammalian host cells reveal that STM is well-adapted to life in specific PCV, referred to as *Salmonella*-containing vacuole, or SCV. The SCV possesses canonical markers of late endosomal compartments, yet allows STM survival and proliferation. The manipulation of the host cell endosomal system mainly mediated by effector proteins of the SPI2-encoded T3SS is central to SCV formation and maintenance (Jennings et al., 2017).

In addition to survival and proliferation in the SCV, further intracellular fates of STM are observed. If the integrity of the SCV is not maintained, STM is exposed to host cell cytosol. This may evoke xenophagic clearance, induce pyroptotic cell death, or may lead to cytosolic hyper-replication resulting in release of highly infected enterocytes as observed for STM (Brumell et al., 2002; Knodler et al., 2010).

In an approach to understand specific virulence mechanisms of TS, a comprehensive comparative transcriptional analyses of intracellular STM and SPA was performed (Cohen et al., 2022). The data revealed various differences in expression of metabolic functions and distinct patterns of expression of flagella genes. We followed potential phenotypic consequences of the distinct expression patterns and observed that a subpopulation of intracellular SPA expresses flagella and is motile in host cell cytosol. Such flagella-mediated cytosolic motility was not observed for STM present in host cell cytosol, and clearly is distinct from intracellular motility evolved by other pathogens, where host cell actin polymerization is hijacked by various surface proteins.

Here we set out to analyze why SPA is released into cytosol, and how host cells respond to intracellular motile SPA. In contrast to intracellular motility mediated by actin polymerization (reviewed in Dowd et al., 2021), intracellular motility of SPA did not allow intercellular spread.

However, flagella-mediated motility enables SPA to avoid xenophagic clearance, as prerequisite of host cell exit of invasion-primed motile SPA.

## Results

### Early escape to host cell cytosol depends on invasion mechanism of SPA

Although STM and SPA belong to the same species, the disease they cause in humans is very different ranging from a self-limiting gastroenteritis in case of STM infection, to severe systemic disease with potentially lethal outcome caused by SPA. We recently compared the intracellular gene expression profiles of STM and SPA during infection and identified differences in the expression of flagella/chemotaxis, SPI1, and carbon utilization pathways, and demonstrated flagella-mediated movement of SPA in cytosol of host cell (Cohen et al., 2022). Flagella-mediated motility was not observed for intracellular STM, despite a small proportion of STM also enters host cell cytosol.

To further investigate phenotypic heterogeneity of intracellular SPA, we performed live cell imaging (LCI) of infected LAMP1-GFP expressing HeLa cells, and compared SPA WT to isogenic mutant strains deficient in flagella synthesis (Δ*fliC*), torque generation (Δ*motAB*), or certain effector proteins of SPI1-T3SS (Δ*sopE*, Δ*sopE2*) or SPI2-T3SS (Δ*sifA*) crucial for early and late SCV integrity. Furthermore, a SPA strain with a synthetic zipper invasion mechanism was investigated, i.e. SPA Δ*invA* [P_*invF*_::*Y.p. inv*] with defective SPI1-T3SS and *Yersinia pseudotuberculosis* (*Y.p.*) Invasin protein Inv synthesized under control of SPI1 promoter P_*invF*_. We quantified the intracellular phenotypes in categories of ‘cytosolic motile’ and ‘cytosolic non-motile’, while ‘other’ includes SPA residing in an SCV or showing cytosolic hyper-replication (**Fig. 1**). In line with previous findings (Cohen et al., 2022), in about 38% of SPA WT-infected host cells harbored cytosolic SPA exhibiting intracellular motility, while in 5% of infected cells cytosolic SPA WT were non-motile (**Fig. 1AB**). SPA mutant strains with defects in maintaining SCV integrity (Δ*sifA*), or lacking SPI1-T3SS effectors (Δ*sopE*, Δ*sopE2*) showed similar results of phenotypes with cytosolic motile SPA Δ*sopE*, Δ*sopE2,* or Δ*sifA* in 30%, 39%, or 40 % of infected cells, respectively. This population was absent in cells infected with non-motile mutant strains lacking the flagella filament subunit FliC (**Fig. 1A** and **Fig. 1C**), or defective in energizing flagella rotation (Δ*motAB*), similar to cells exposed to the antibiotics cefotaxime and ciprofloxacin affecting bacterial cell wall synthesis and DNA replication during infection. Because Δ*motAB* and Δ*fliC* strains are non-motile, we also tested if motility during invasion may affect intracellular phenotypes. For this, expression of *motAB* was placed under control of promoter P_*tetA*_, and induced by addition of anhydrotetracycline (AHT) (Schulte et al., 2019). We induced expression of *motAB* with AHT in bacterial culture and omitted AHT during infection and the following incubation time to avert motility in infected HeLa cells. SPA Δ*motAB* [P_*tetA*_::*motAB*] showed increased invasion compared to Δ*motAB* (**Fig. S 4**), but also completely lacked the cytosolic motile subpopulation. Further, we set out to evaluate the contribution of the mode of invasion to cytosolic release and subsequent intracellular motility using mutant strain Δ*invA* incapable of translocation of trigger invasion-mediating SPI1-T3SS effectors. We introduced a plasmid for expression of *Y.p. inv* under control of SPI1 promoter P_*invF*_, allowing expression under similar conditions as the SPI1-T3SS, and therefore conferring zipper invasion to SPA. Interestingly, for this strain the subpopulation of cytosolic motile bacteria was highly reduced to 1% of infected cells, the cytosolic subpopulation was almost absent (5%), and SPA predominantly residing in SCV (**Fig. 1A** and **Fig. 1D**). These data confirm previously data stating trigger invasion as driving force for cytosolic release (Röder & Hensel, 2020).

**Fig. 1.**
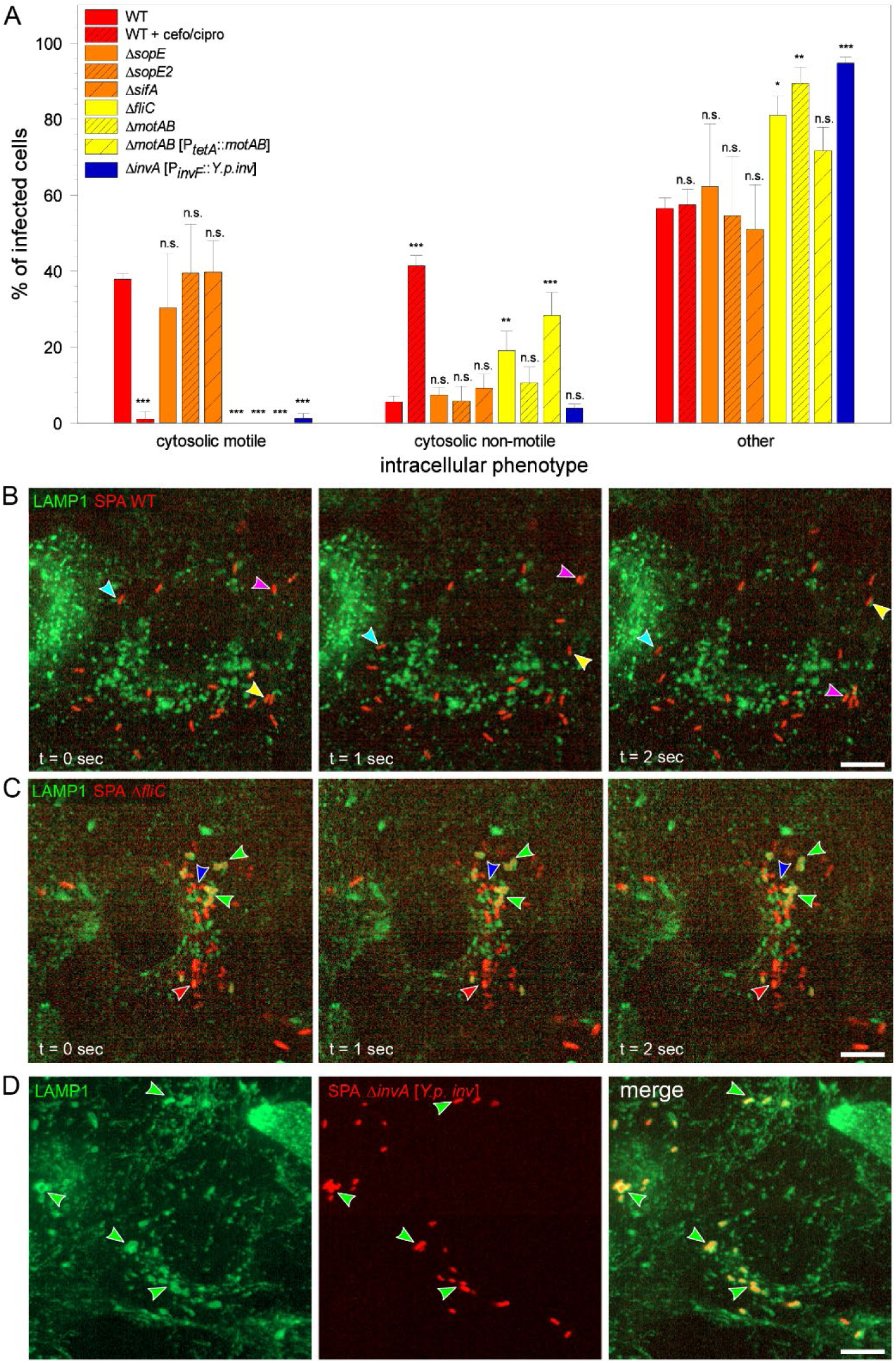
Quantification of intracellular SPA phenotypes. **A-E.** HeLa cells stably expressing LAMP1-GFP (green) were infected with *Salmonella* Paratyphi A (SPA) WT or isogenic mutant strains expressing mCherry (red) at MOI 30, 60, or 90 for WT, Δ*invA* [P_*invF*_::*Y.p. inv*], or Δ*fliC*, Δ*motAB*, respectively. **A.** Quantification of intracellular phenotypes was performed during live cell imaging. From 4-6 h p.i., at least 100 infected cells per strain were examined for intracellular phenotypes. Data are means and standard deviations (SD) from three independent experiments. Statistical analysis was performed with one-way ANOVA and is indicated as n.s., not significant; *, p < 0.05; **, p < 0.01; ***, p < 0.001. **B, C.** Time-lapse images of infected HeLa-LAMP1-GFP cells showing movement of three cytosolic SPA WT (**B**, turquoise, magenta and yellow arrowhead), no movement of cytosolic SPA Δ*fliC* (**C**, red and blue arrowhead) and SPA Δ*fliC* residing in SCV (**C**, green arrowheads). **D.** Maximum intensity projection of HeLa-LAMP1-GFP cells infected with SPA Δ*invA* [P_*invF*_::*Y.p. inv*] residing in SCV (green arrowheads). Scale bars: 10 µm.

### Cooperative trigger invasion of SPA leads to increased membrane damage of SCV

It was previously described that the mode of trigger invasion and also strain-specific equipment with SPI1-T3SS effectors affect cytosolic release following host cell entry and that trigger invasion is often cooperative between *Salmonella* (Lorkowski et al., 2014; Röder & Hensel, 2020). We set out to investigate membrane ruffling during invasion by LCI of different STM serovars and SPA WT as well as SPA invading through zipper mechanism (**Fig. 2**, **Fig. S 1**, **Movie 1, Movie 2, Movie 3, Movie 4**). We observed rather small membrane ruffling for STM NCTC 12023 (**Fig. 2A**, **Fig. S 1, Movie 1**) and more extensive membrane ruffling for STM SL1344 (**Fig. 2B, Fig. S 1, Movie 2**). Prolonged and extensive membrane ruffling of a larger cell area was observed for SPA WT (**Fig. 2C, Fig. S 1, Movie 3**). SPA Δ*invA* [P_*invF*_ *Y.p. inv*] showed lowest actin rearrangement at the invasion site (**Fig. 2D, Fig. S 1, Movie 4**). Image analysis of foci of invasion indicated that STM NCTC 12023 entry occurs mostly by single bacteria at invasion sites (mean = 1.6 STM cells per site), whereas during the invasion of STM SL1344 multiple bacteria were involved in the process (mean = 3.2 STM cells per site). SPA WT showed highest accumulation of bacteria at sites of membrane rearrangement (mean = 7.3 SPA cells per site), but numbers were far lower when SPA invaded through zipper mechanism (mean = 2.1 SPA cells per site).

**Fig. 2.**
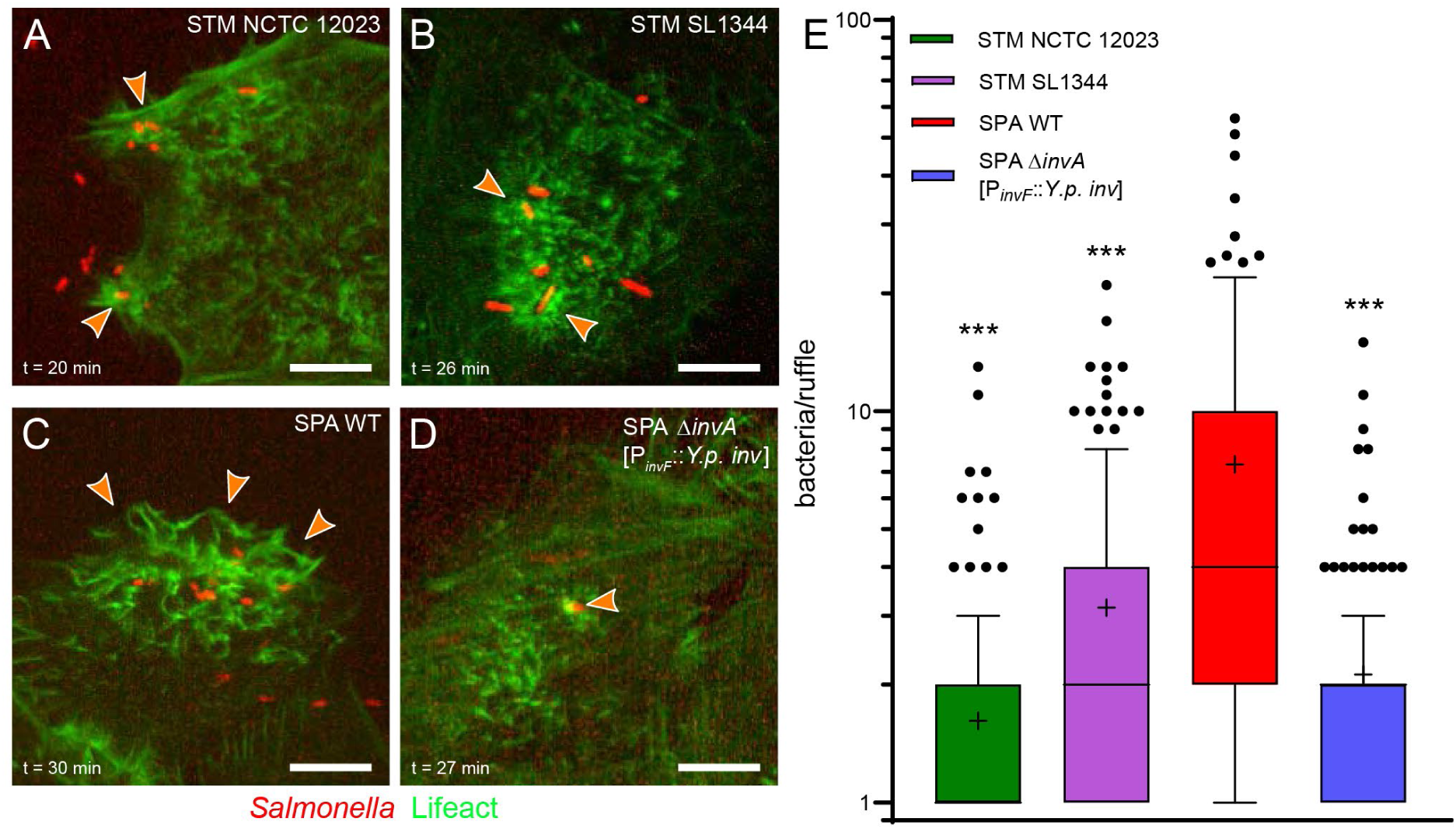
SPA induces massive membrane ruffles during trigger invasion, and invades in highly cooperative manner. **A-D**. HeLa cells stably expressing Lifeact-eGFP (green) were infected with STM NCTC 12023 (**A**), STM SL1344 (**B**), SPA WT (**C**), or SPA Δ*invA* [P_*invF*_::*Y.p. inv*] (**D**) expressing mCherry (red) at MOI 75. Cells in chambered coverslips were infected on microscope stage and imaged for 1 h by spinning disc confocal microscopy with intervals of 3-5 min between images. Still images from time-lapse are shown. Arrowheads indicate sites of ongoing invasion. Scale bars: 10 µm. **E.** Image series were assessed for numbers of invading bacteria per ruffle. Shown are Tukeýs box plots of invading bacteria per ruffle during infection with error bars including data of 1.5 x interquartile range (IQR). Outliers beyond the 1.5 x IQR are shown as dots. Middle lines and “+” denote median and mean value, respectively. Compiled from three independent experiments, 200, 169, 170, and 130 individual invasion events were analyzed for STM NCTC 12023, STM SL1344, SPA WT, and SPA Δ*invA* [P_*invF*_::*Y.p. inv*], respectively,. Statistical analysis compared to SPA WT was performed with unpaired two-tailed t test and is indicated as *** for p < 0.001.

### Trigger invasion enhances membrane damage at nascent SCVs

Previous work revealed that the SCV is prone to rupture, and damages compartments are targeted by membrane damage sensors and repair mechanisms such as galectins, sphingomyelinases, or the ESCRT machinery (Ellison et al., 2020; Göser et al., 2020; Paz et al., 2010; Thurston et al., 2012). We used an engineered version of equinatoxin II (EqtSM) as rapid membrane damage reporter that is binding cytosol-exposed sphingomyelin in damages endosomal membranes (Deng et al., 2016; Niekamp et al., 2022). HeLa cells expressing Halo-tagged EqtSM were infected by SPA WT or SPA Δ*invA* [P_*invF*_::*Y.p. inv*] to assess contribution of invasion mechanism to membrane damage of the nascent SCV (**Fig 3**). LCI revealed that trigger invasion by SPA WT led to 45% of SCV positive for EqtSM indicating transient membrane damage, while only 14% of SCV were EqtSM-positive if SPA invaded through zipper mechanism (**Fig 3D**). The results underline the critical role of entry mechanism for SCV integrity, and for subsequent intracellular lifestyle of *Salmonella*.

**Fig 3.**
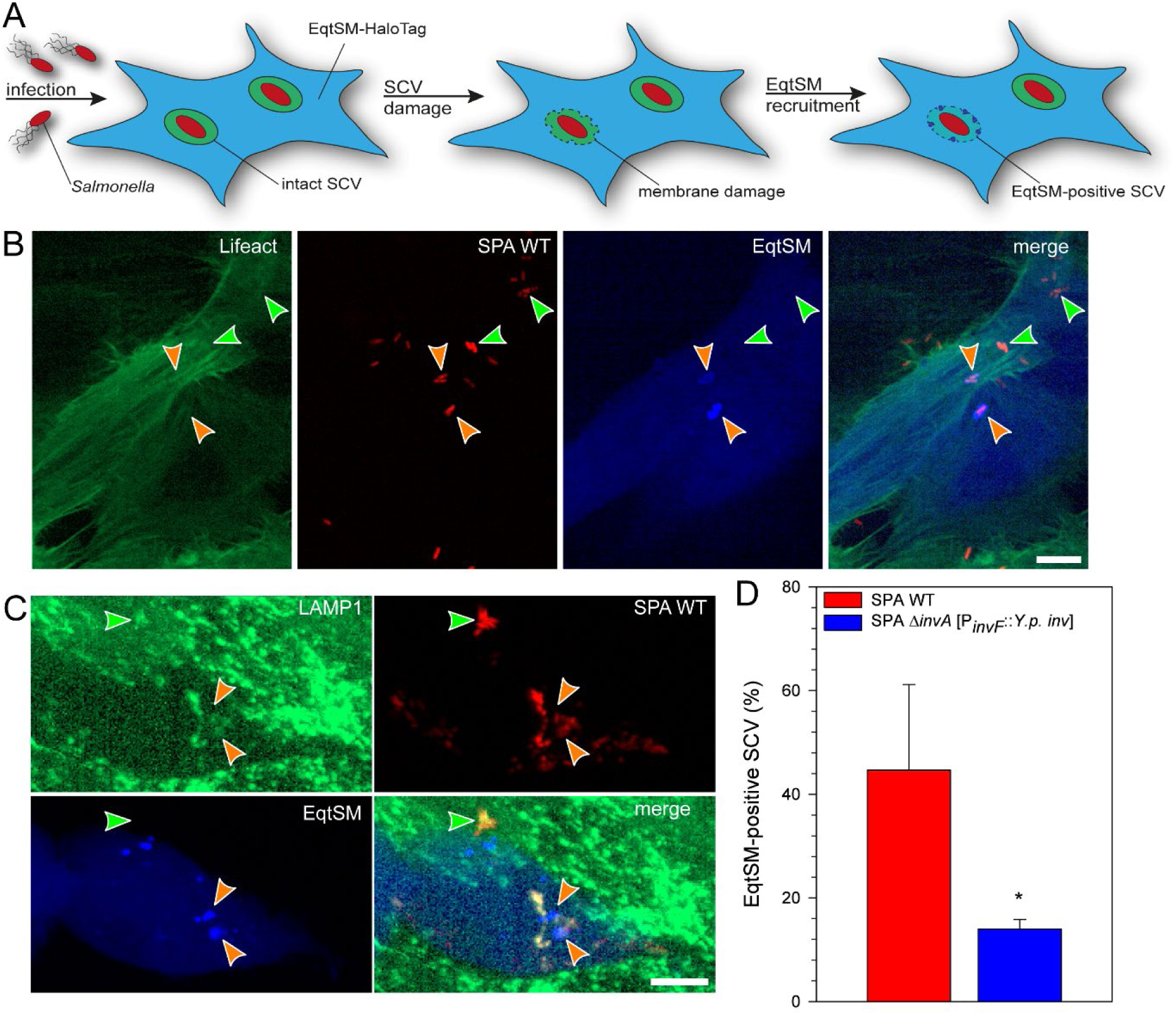
Trigger invasion by SPA destabilizes nascent SCVs. **A.** To localize sites of SPA-induced membrane damage and to quantify the frequency of SCV damage, HeLa cells stably expressing LAMP1-GFP (**B**) or Lifeact-eGFP (**C**) (green) were transfected for expression of EqtSM-HaloTag and labeled with Janelia Fluor 646 HaloTag ligand (blue). Subsequently, cells were infected with SPA WT or SPA Δ*invA* [P_*invF*_::*Y.p. inv*] expressing mCherry (red). Infected cells were imaged 1-2 h p.i. and representative infected cells are shown as maximum intensity projection. Arrowheads indicate intact SCVs (green), or SCVs targeted by EqtSM (orange). Scale bars: 10 µm. **D**. SCVs were scored for association with EqtSM-HaloTag. Data are means and SD from three independent experiments. Statistical analysis was performed with unpaired two-tailed t test and is indicated as * for p < 0.05.

### Motility of cytosolic SPA interferes with xenophagic clearance

A subset of intracellular pathogens such as *Listeria monocytogenes* (*L.m*.) or *Shigella flexneri* actively escape the early PCV. As the host cell cytosol is a hostile environment for cytosolic bacteria, e.g. due to recognition and clearance by cell-autonomous defense mechanism such as the autophagosomal machinery, these pathogens evolved sophisticated defense mechanisms to avoid decoration and degradation by expression of proteins interfering and inhibiting xenophagy. Such mechanisms have not been reported for cytosolic *Salmonella*. An event preceding autophagic clearance of cytosolic bacteria is ubiquitylation of their surface structures, such as LPS (Otten et al., 2021). We also observed ubiquitylation of SPA located in host cell cytosol (**Fig. S 2**), and analyzed if subsequent xenophagic clearance occurs. Alternatively, we hypothesized that cytosolic motility of SPA may possibly interfere with xenophagy. To investigate on single cell level the capture of SPA by autophagosomal membranes, we deployed late-autophagosomal membrane protein LC3B as marker, generated a HeLa line stably expressing LC3B-GFP, and used this cell line for infection by SPA and LCI. Individual intracellular SPA were scored for motility and association with LC3B-positive membranes (**Fig. 4**, **Movie 5**, **Movie 6**). We compared SPA WT to SPA Δ*motAB* [P_*tetA*_::*motAB*] (motile during infection, but no intracellular motility), and SPA WT treated with cefotaxime and ciprofloxacin (non-motile and non-replicative) regarding levels of LC3B decoration. SPA-infected cells showed heterogenous populations as representative images in **Fig. 4D-F** illustrate. The number of bacteria and also the degree of LC3B decoration varied from cell to cell (**Fig. 4D-F**, **Movie 5**, **Movie 6**).

**Fig. 4.**
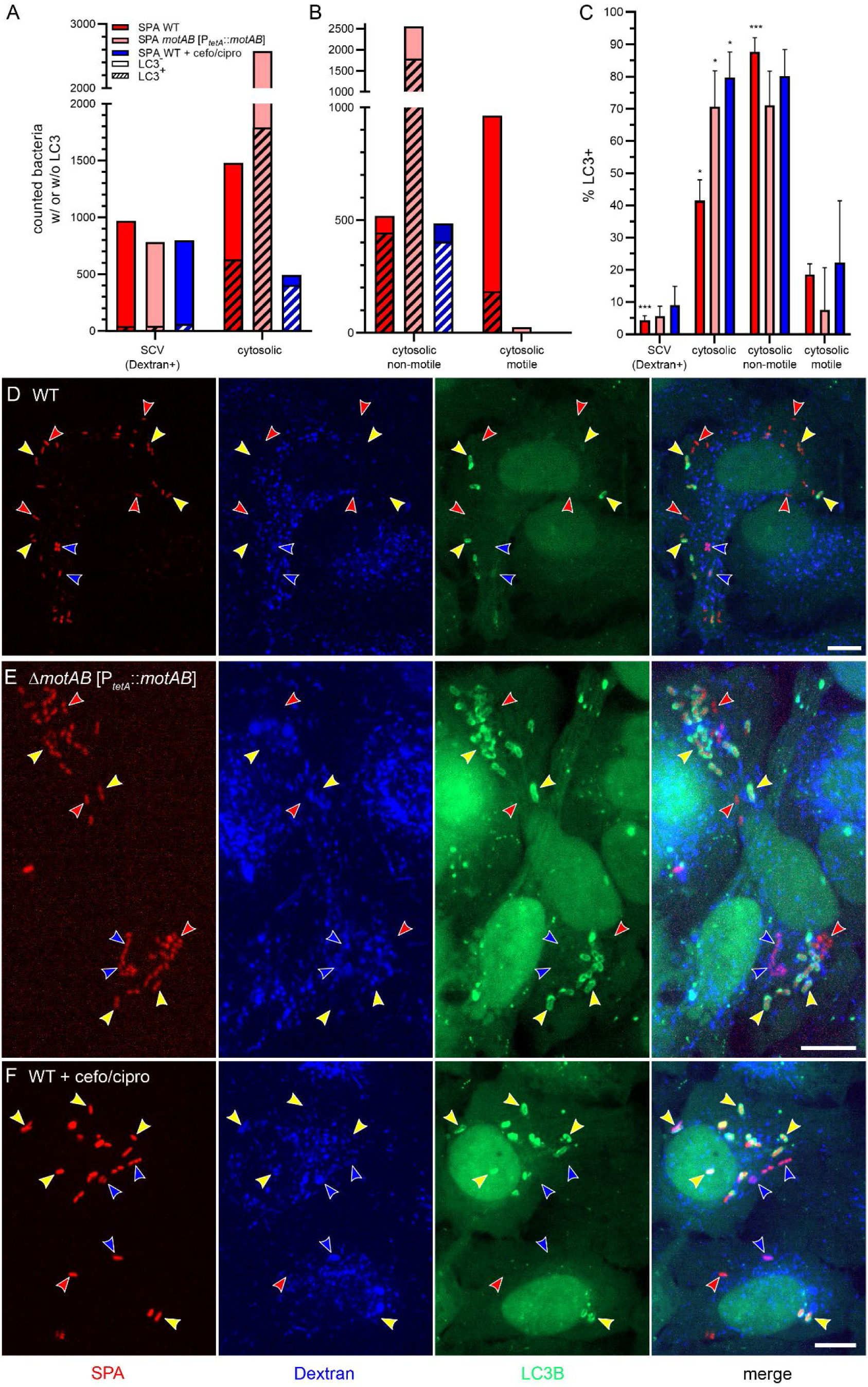
Cytosolic non-motile SPA are targeted by autophagosomal membranes. **A-C**. HeLa cells stably expressing LC3B-GFP (green) were pulse-chased with Dextran-Alexa647 (blue) and infected with SPA WT or SPA Δ*motAB* [P_*tetA*_::*motAB*] expressing mCherry (red) at MOI 30. 100 µg x ml^-1^ cefotaxime and ciprofloxacin were added to infected cells at 2 h p.i. if indicated. At 3-4 h p.i., randomly selected cells were imaged and later assessed for SPA associated with LC3B-positive membranes. **A, B.** Total counts of SPA with (dashed bars) or without (bars) LC3B decoration, compiled from three independent experiments. **B.** Cytosolic SPA quantified in **A** were further scored as motile or non-motile, and assessed for LC3B decoration. **C.** Percentage of SPA targeted by LC3B-containing membranes calculated from data shown in **A** and **B**. Data are means and SD of 195 infected cells with 2,450 bacteria for WT, 171 infected cells with 3,360 bacteria for Δ*motAB* [P_*tetA*_::*motAB*], and 215 infected cells with 1,291 bacteria for WT + cefo/cipro from three independent experiments. Statistical analysis was performed with one-way ANOVA and is indicated as n.s., not significant; *, p < 0.05; **, p < 0.01; ***, p < 0.001. **D-F.** Representative images showing subpopulations in SPA-infected cells. Arrowheads indicate SPA residing in SCV (blue), cytosolic SPA (red), LC3B-decorated SPA (yellow). Images are shown as single Z-plane from stack (**D**) or as maximum intensity projection (**E, F**). Scale bars: 10 µm.

Image analysis of more than 170 infected cells with over 1,200 bacteria per condition (**Fig. 4A**, **Fig. 4B**) revealed that residing in SCV efficiently avoids LC3B decoration for all strains (4.5% LC3B-positive SPA WT, 5.7% LC3B-positive SPA Δ*motAB* [P_*tetA*_::*motAB*]), with the highest rate of LC3B decoration observed in cefo/cipro-treated SPA WT (9% LC3B-positive, **Fig. 4C**), possibly due to failure of maintaining SCV integrity due to antibiotics treatment. However, the cytosolic populations of the respective strains showed significant differences regarding LC3B decoration. The enteric cytosolic population of SPA WT showed about 41.5% LC3B-positive bacteria, whereas cytosolic populations of SPA Δ*motAB* [P_*tetA*_::*motAB*], and SPA WT + cefo/cipro both showed a high frequency of LC3B decoration (70.6%, 79.7%, respectively, **Fig. 4C**). We next dissected the cytosolic population into groups of non-motile SPA and motile SPA to analyze effects of motility on xenophagic escape. Surprisingly, SPA WT showed a high frequency of LC3B decoration for the non-motile subpopulation (87.7%), but low frequency for the motile subpopulation (18.6%). In contrast, the motile fraction was almost absent for SPA Δ*motAB* [P_*tetA*_::*motAB*] and SPA WT + cefo/cipro strains with only 31 and 8 counted bacteria, respectively (**Fig. 4B**). Thus, the frequency of LC3B decoration of the cytosolic population of these strains reflects that of the non-motile fraction with 71.1% and 80.2%, respectively. We conclude that cytosolic motility is a major factor for xenophagic escape of SPA.

To further investigate the heterogenous fate of individual intracellular SPA subpopulations, correlative light and electron microscopy (CLEM) analyses were performed to reveal ultrastructure and membrane association (**Fig. 5**). Light microscopy allowed identification of cytosolic motile and non-motile, as well as SCV-bound SPA for subsequent analyses by transmission electron microscopy (TEM). Whereas cytosolic motile SPA showed no contact to membranous structures, we observed SCV harboring SPA with distinct single membrane compartment in close contact to the bacterial envelope (**Fig. 5D, E**). Cytosolic non-motile SPA showed association with LC3B-positive membranes (**Fig. 5F**). The LC3B signal correlated double membrane structures characteristic for autophagosomes. We also detected SPA that were partially enclosed by LC3B-positive membranes, indicating either ongoing autophagosomal capture, or futile xenophagy.

**Fig. 5.**
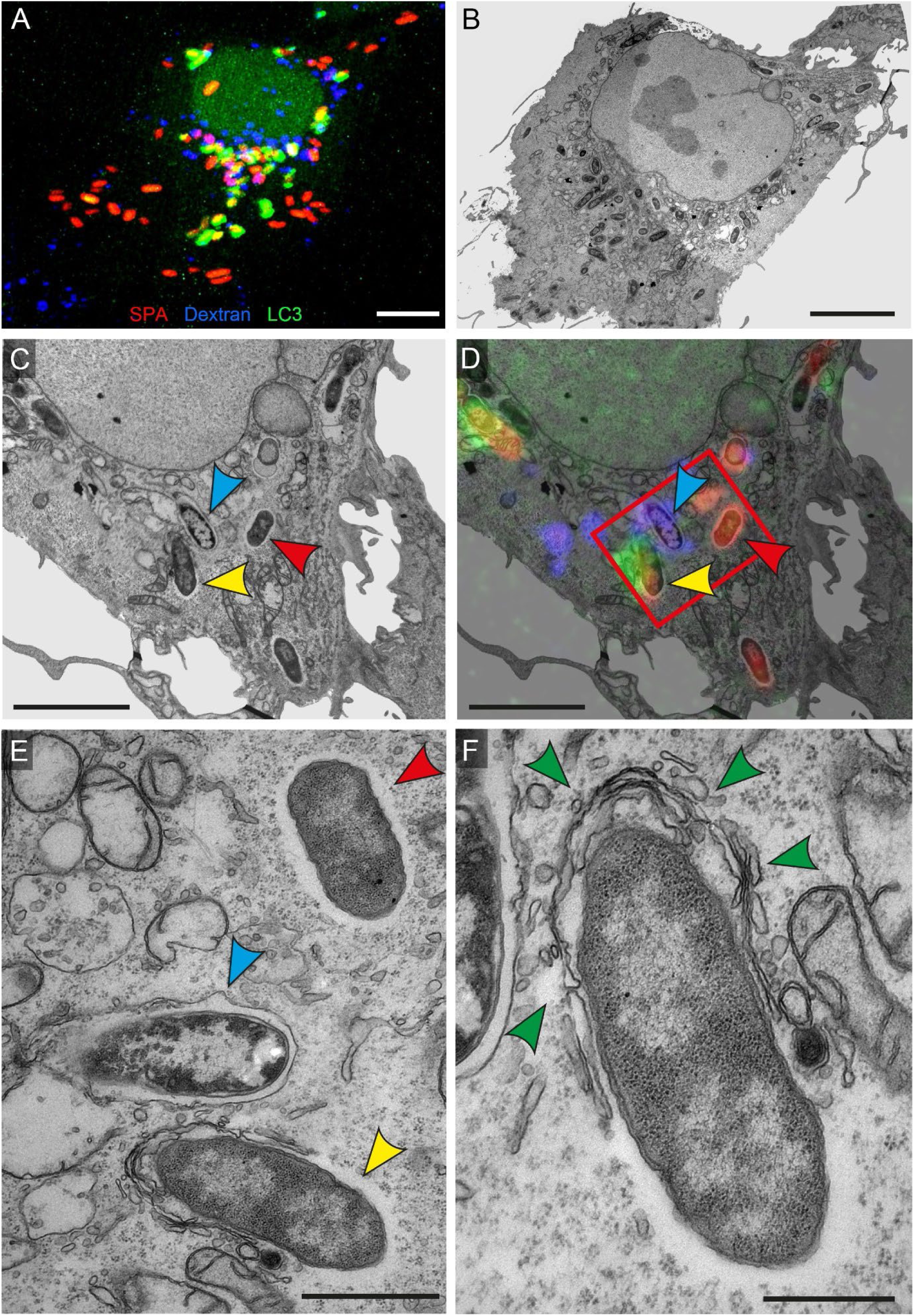
Various degrees of xenophagic capture of distinct subpopulations of intracellular SPA. HeLa cells expressing LC3B-GFP (green) were pulse-chased with Dextran-Alexa647 (blue) and infected with SPA WT expressing mCherry (red) at MOI 30. Cells were fixed 4 h p.i. during LCI (**A**), and subsequently processed for transmission electron microscopy (TEM) of ultrathin sections. LCI and TEM images were superimposed for correlation (**D**) and details are shown at higher magnification (**E**, **F**). Arrowheads indicate SPA residing in SCV (blue), cytosolic SPA (red) and SPA targeted by LC3B-positive autophagosomal membrane (yellow). Partial enclosure by autophagosomal membranes is indicated by green arrowheads. Scale bars: 10 µm (**A**, **B**), 5 µm (**C**, **D**), 1 µm (**E**), 500 nm (**F**).

### Increasing LC3B decoration hinders motility of cytosolic SPA

For many bacteria of various cell shapes, flagellar rotation provides force for locomotion in their habitat. The rotation of one or multiple flagellar filaments is thereby mediating phases of straight swimming, alternating with phases of tumbling allowing orientation in chemical gradients. We set out to investigate the maximum velocity of flagella-mediated movement of cytosolic SPA. As previously demonstrated for multiple other bacteria, swimming speed is dependent on type of flagellation and rotation speed of the flagellum (reviewed in Wadhwa & Berg, 2022). First, we compared maximum velocity during different growth conditions in LB medium (**Fig. 6A**). SPA cultures grown under aerobic conditions showed a median maximum velocity of 11 µm s^-1^ when grown overnight and 42 µm s^-1^ when grown for 3.5 h while SPI1-inducing microaerophilic conditions led to a median maximum velocity of 26 µm s^-1^. Other intracellular pathogens such as *L.m*. utilize actin polymerization to energize locomotion in host cell cytosol. We observed median maximum velocity of cytosolic *L.m*. of 0.5 µm s^-1^ (**Fig. 6B**) which was within the range of previously published data (Lacayo & Theriot, 2004). In contrast, cytosolic SPA were able to move almost as fast as SPA grown in media under microaerophilic conditions (25 µm s^-1^; **Fig. 6C**).

**Fig. 6.**
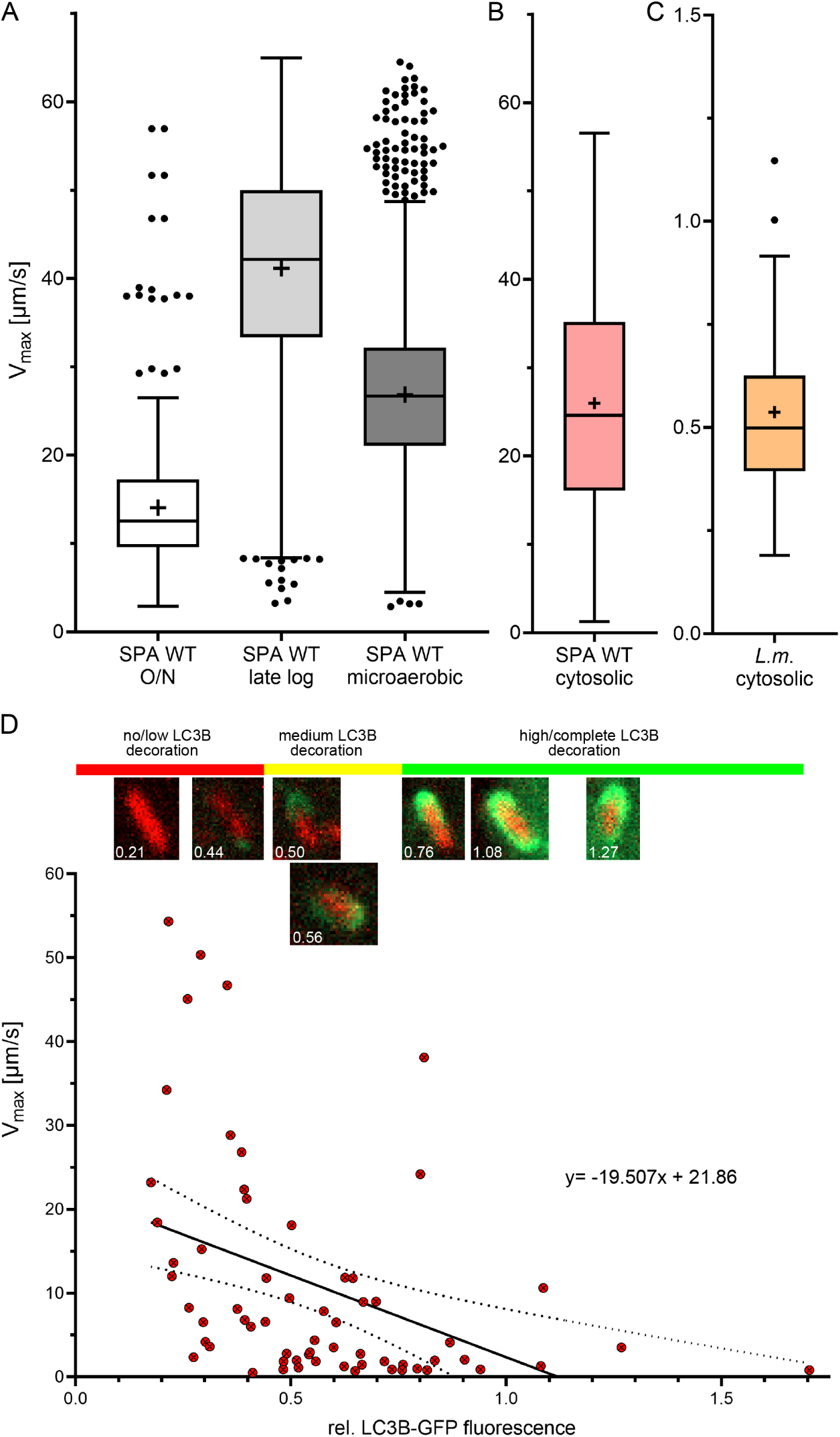
Xenophagic capture reduces velocity of intracellular motile SPA. Single bacterial cells after culture in broth media (**A**), or within infected host cells (**B**, **C**) were registered by LCI. Subsequently, bacterial motility was analyzed by automated (**A**, **C**) or manual (**B**) tracking analyses. **A-C.** Maximum velocity of single bacterial cells from tracking SPA grown in culture (**A**), or intracellular SPA (**B**) or *L.m.* (**C**). For single cell analyses, tracks of 476 SPA WT from O/N culture, 7,058 SPA WT from late log culture, 2,998 SPA WT from micro-aerobic culture were collected from one experiment. For intracellular bacteria, tracks of 16,248 cytosolic SPA WT were collected from three independent experiments, and 42 tracks for *L.m.* from one experiment. Data are shown as Tukeýs box plots with error bars including data of 1.5 x IQR. Outliers beyond the 1.5 x IQR are shown as dots. Middle line denotes median and “+” mean value. **D.** Plotting of SPA-associated relative LC3B-GFP intensity against maximum velocity of individual cytosolic SPA. Representative images from measurement of fluorescence intensity are shown as single Z-plane from image stacks. Linear regression and 95% confidence bands were calculated for analyses of 63 cytosolic SPA from two independent experiments.

As described above, a subpopulation of LC3B-decorated SPA was still able of movement (**Fig. 4**, **Movie 5**, **Movie 6**). We hypothesized that increased autophagosomal capture delimits intracellular motility of SPA, or in turn, intracellular motility prevents autophagosomal capture. To test these possible correlations, we determined the signal intensity of SPA-associated LC3B-GFP as proxy of degree of autophagosomal capture of individual SPA, and determined by tracking of the same cytosolic SPA cells the maximum velocity (Fig. 6D, Fig. S 3). Plotting LC3B-GFP signal intensity vs. maximal velocity of individual SPA indicated that velocity decreases with increasing association with autophagosomal membranes. Linear regression with principal component analysis showed a negative slope (y=-19.5x + 21.9) with maximum velocity, dropping to <5 µm s^-1^ at a relative LC3B-GFP fluorescence of 0.86 which reflects an almost complete enclosure of SPA. We conclude that flagella-mediated cytosolic motility of SPA counteracts host cell xenophagy.

## Discussion

Our study investigated the causes of the recently described cytosolic motility of SPA (Cohen et al., 2022) and its contributions to intracellular lifestyle. We demonstrated that mode of invasion and equipment with SPI1-T3SS effector proteins is substantial for membrane damage at the nascent SCV membrane and cytosolic release in early stage of infection. Furthermore, we showed that non-motile SPA are targeted by autophagosomal membranes, However, the population of cytosolic motile SPA is able to delay or escape xenophagic clearance as infection progresses. Our work adds flagella-mediated motility of intracellular bacteria to the mechanisms of evasion of host cell xenophagy.

The main findings of this study are summarized in **Fig 7**. We propose that release of SPA into cytosol occurs early after invasion, and is likely due to the inability to generate a stable nascent SCV (**Fig 7E**). We frequently observed clusters of invading SPA that induce high levels of local actin recruitment, and extensive membrane ruffles. Such events likely lead to incomplete SCV formation, and may allow clusters of bacteria to directly enter cytosol. This early exit from a labile nascent SCV occurs so rapid that the transcriptional profile cannot be adjusted to the SCV-specific repression of SPI1 and flagella regulons. We propose that a large proportion of intracellular SPA maintains the transcriptional profile of invading bacteria. This hypothesis may be tested by labelling of flagella, infection, and live cell imaging to follow the fate of flagella on SPA single cells after invasion.

**Fig 7.**
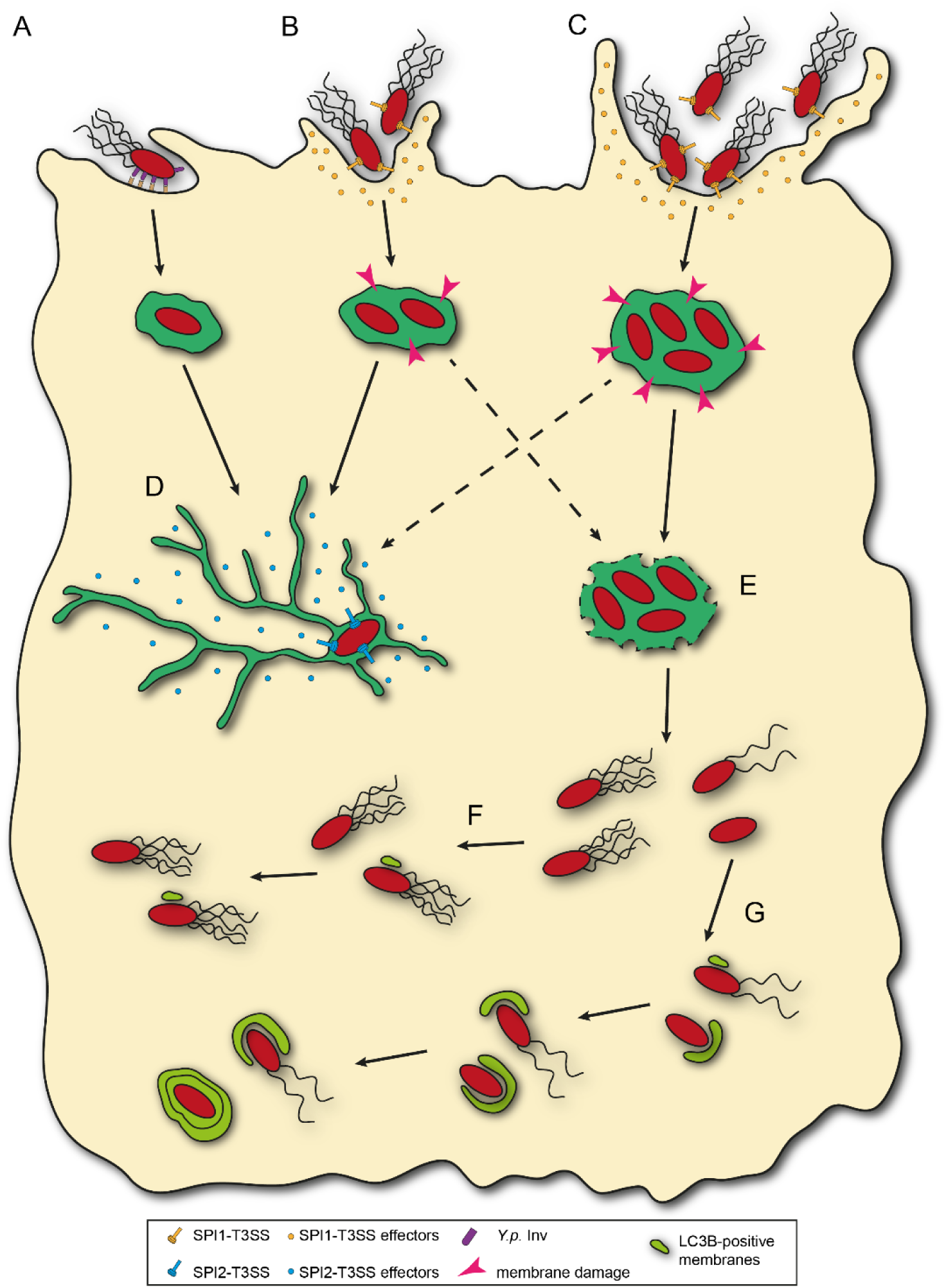
Model for SPA entry into host cell cytosol and autophagic decoration. Zipper or trigger invasion with limiting extend of membrane ruffling (**A**, **B**) primarily lead to an intracellular lifestyle with SCV maturation (**D**). Trigger invasion with extensive membrane ruffling and increased cooperative invasion (**C**) lead to membrane damage and rupture of the nascent SCV (**E**) and a cytosolic lifestyle of SPA. Flagellated and motile cytosolic SPA delay or escape xenophagic capture (**F**). Non-motile cytosolic SPA are captured by LC3B-positive membranes, while flagella-mediated motility permits evasion of xenophagy (**G**).

SPA possesses both SopE and SopE2. We previously demonstrated that STM harboring SopE2 and SopE induce more severe membrane ruffles compared to the majority of STM strains only harboring SopE2 (Röder & Hensel, 2020). The more overt trigger invasion by SopE-positive STM was associated with increased cytosolic release, also occurring early after invasion. This may be similar to damage of nascent SCV induced by SPA. However, we did not observe flagella-mediated intracellular motility of SopE/SopE2-positive STM SL1344 (Cohen et al., 2022), indicating further differences between STM and SPA. SPA possesses functional *sopE* and *sopE2*, while STY strains possess functional *sopE* and pseudogenic *sopE2* (Valenzuela et al., 2015), thus damage of nascent SCV after invasion of STY may be less pronounced compared to SPA.

A further factor possible contributing to decreased stability of the nascent SCV is the altered function of SptP, a SPI1-T3SS effector protein with activity antagonistic to SopE. Prior work showed that host cells fail to restore normal actin cytoskeleton organization when SptP is defective and SopE activity overrules (Kubori & Galan, 2003). Recent work by Johnson *et al*. (Johnson et al., 2017) demonstrated that SptP in STY and SPA have several AA exchanges that affect the binding of chaperone SicP, and by this the efficiency of translocation.

Virtually all intracellular bacterial or parasitic pathogens adapted to cytosolic lifestyle have convergently evolved mechanisms to recruit host cell G-actin for polar polymerization to F-actin, resulting in actin-based motility (ABM) (reviewed in Dowd et al., 2021). ABM enables cell-to-cell spread, and thus enables infection of new host cells without demanding exit and exposure to antimicrobial mechanisms acting on extracellular bacteria. We did not observe intercellular spread of SPA. While ABM is slower compared to flagella-mediated cytosolic motility of SPA, the higher force generated by actin polymerization leads to generation of membrane protrusions and ultimately to intercellular spread. Furthermore, switching between swimming and tumbling motility restricts generation of membrane protrusions that are productive in infection of neighboring cells. Yet, we have to consider that the applied model of confluently cultured non-polarized epithelial cells may not reflect host cell types relevant during infection of human hosts as main host cells of SPA are phagocytic cells, leading to dissemination through the lymphatic system and further colonization of internal organs such as the liver, spleen, bone marrow and gall bladder. Motility of SPA inside phagocytic cell line U937 has only been observed in spacious vacuoles rather than in the cytosol of host cells (Cohen et al., 2022), probably because presence of bacteria in cytosol of phagocytic cells most likely results in inflammatory cell death, so-called pyroptosis (Castanheira & Garcia-Del Portillo, 2017). Similar observations were made for STM in gallbladder epithelium (Knodler et al., 2010). In contrast to STM, SPA shows a lower replication rate in host cells (Reuter et al., 2021). Defects in metabolic utilization of nutrients in host cell cytosol such as glucose-6-phosphate and amino acids, and low SPI2-mediated replication inside SCV may contribute to reduced replication numbers of intracellular SPA (Cohen et al., 2022; Forest et al., 2010; Holt et al., 2009).

The cytosolic presence of pathogens such as *L. monocytogenes* or *S. flexneri* induce host cell-intrinsic defense mechanisms, resulting in ubiquitination and xenophagy. Xenophagy is partially avoided by ABM (Yoshikawa et al., 2009), but additional mechanisms appear required to efficiently avoid autophagosomal degradation. These include the T3SS effector protein IcsB of *S. flexneri* (Ogawa et al., 2005) with an acyltransferase activity affecting small GTPases and membrane fusion machinery (Liu et al., 2018), or secreted Listeriolysin O and phospholipases of *L. monocytogenes* targeting autophagosomal membranes (Birmingham et al., 2007). For SPA, the high speed of flagella-mediated motility is sufficient to enable protection of a subpopulation of cytosolic SPA against xenophagy (**Fig 7F**). Xenophagic clearance of cytosolic SPA is likely an arms race between host cell and pathogen. If the autophagosomal machinery is rapidly delivered to SPA and its flagella filament, motility is stopped and autophagy can be completed before SPA motility and proliferation initiates (**Fig 7G**). The mode of trigger invasion by clusters of SPA, and resulting simultaneous cytosolic presence of several SPA cells likely contributes to rapid exhaustion of xenophagic capacity of the host cell, and inability to control SPA proliferation.

We showed that the cytosolic population is escaping xenophagic clearance by flagella-based motility. Motile bacteria showed same maximum velocity in host cell cytosol as in culture media, interfering with decoration by autophagosomal membranes, although cytosolic bacteria are already ubiquitylated. If decoration by LC3B-positive membranes is only delayed to late time points of infection and if this also leads to lysosomal degradation of engulfed bacteria deserves further investigation. For several *Yersinia* strains it was reported that degradation by fusion with lysosomes is inhibited and *Yersinia* is able to replicate in a non-acidic autophagosome (Connor et al., 2018; Lemarignier & Pizarro-Cerda, 2020; Moreau et al., 2010; Straley & Harmon, 1984). Further investigations assessing acidification of LC3B-positive compartments and interfering with autophagosome formation using Lysotracker, siRNA approaches, and autophagy inhibitors such as 3-MA in combination with replication assays could help to understand the host-pathogen interactions in SPA in more detail.

During systemic infections by TS induction of massive inflammatory response due to bacteria-induced extrusion of infected cells which and not observed ad this is considered to contribute to stealth strategy of TS (reviewed in Dougan & Baker, 2014; Hiyoshi, Tiffany, et al., 2018). Certain features of immune evasion, such as avoidance of neutrophil oxidative burst have convergently evolved, i.e. by expression of Vi capsule (STY) or the very long O-antigen of LPS (SPA) (Hiyoshi, Wangdi, et al., 2018). Flagella-mediated intracellular motility has not been reported for STY, and we did not observe increased numbers of cytosolic STY, nor detected flagella expressed by intracellular STY. Functional flagella are expressed by intracellular *Legionella pneumophila* during transition from the replicative to non-replicative, transmissive forms, and a recent study showed that flagellated subpopulations emerge in the *Legionella*-containing vacuole (Schell et al., 2016; Striednig et al., 2021). This transition occurs at the end of the intracellular replication cycle, and stimulates host cell pyroptotic death, and release of *L. pneumophila* for new rounds of host cell infection (Schell et al., 2016). We did not observe induction of host cell death by cytosolic motile SPA, but this host cell response is highly cell type dependent. Further understanding of physiological consequences of intracellular motility of SPA demand analyses in improved infection models such as organoids of human origin (Sepe et al., 2020) that are capable to simulate tissues closely related to *in vivo* conditions.

## Acknowledgements

We thank Ohad Gal-Mor (Tel HaShomer, Israel) and the members SalHostTrop consortium (InfectEra 3) for fruitful discussions on intracellular lifestyle of SPA and other TS. We thank Hans-Peter Schmitz (Div. Genetics, UOS) for help with FACS of HeLa-LC3B-GFP cells. *Listeria monocytogenes* expressing RFP was kindly provided by Martin Loessner (ETH Zürich). This work was supported by the DFG through grant HE1964/23-1 as part of priority program SPP 2225 ‘Exit strategies’, and SReferences

## Materials and Methods

### Bacterial strains and growth conditions

In this study, *Salmonella enterica* serovar Typhimurium (STM) NCTC 12023, STM SL1344 and *S. enterica* serovar Paratyphi A (SPA) 45157 were used as wild-type (WT) strains. All mutant strains are isogenic to the respective WT and **Table 1** shows the characteristics. STM and SPA strains were routinely cultured at 37 °C on Luria-Bertani (LB) agar or in LB broth using a roller drum at 60 rpm. *Listeria monocytogenes* was grown on brain heart infusion (BHI) agar or in BHI broth using a roller drum at 60 rpm. Antibiotics for maintenance of plasmids listed in **Table 2** were added to LB in concentrations of 50 µg x ml^-1^ carbenicillin, 50 µg x ml^-1^ kanamycin, or 12 µg x ml^-1^ chloramphenicol.

**Table 1.**
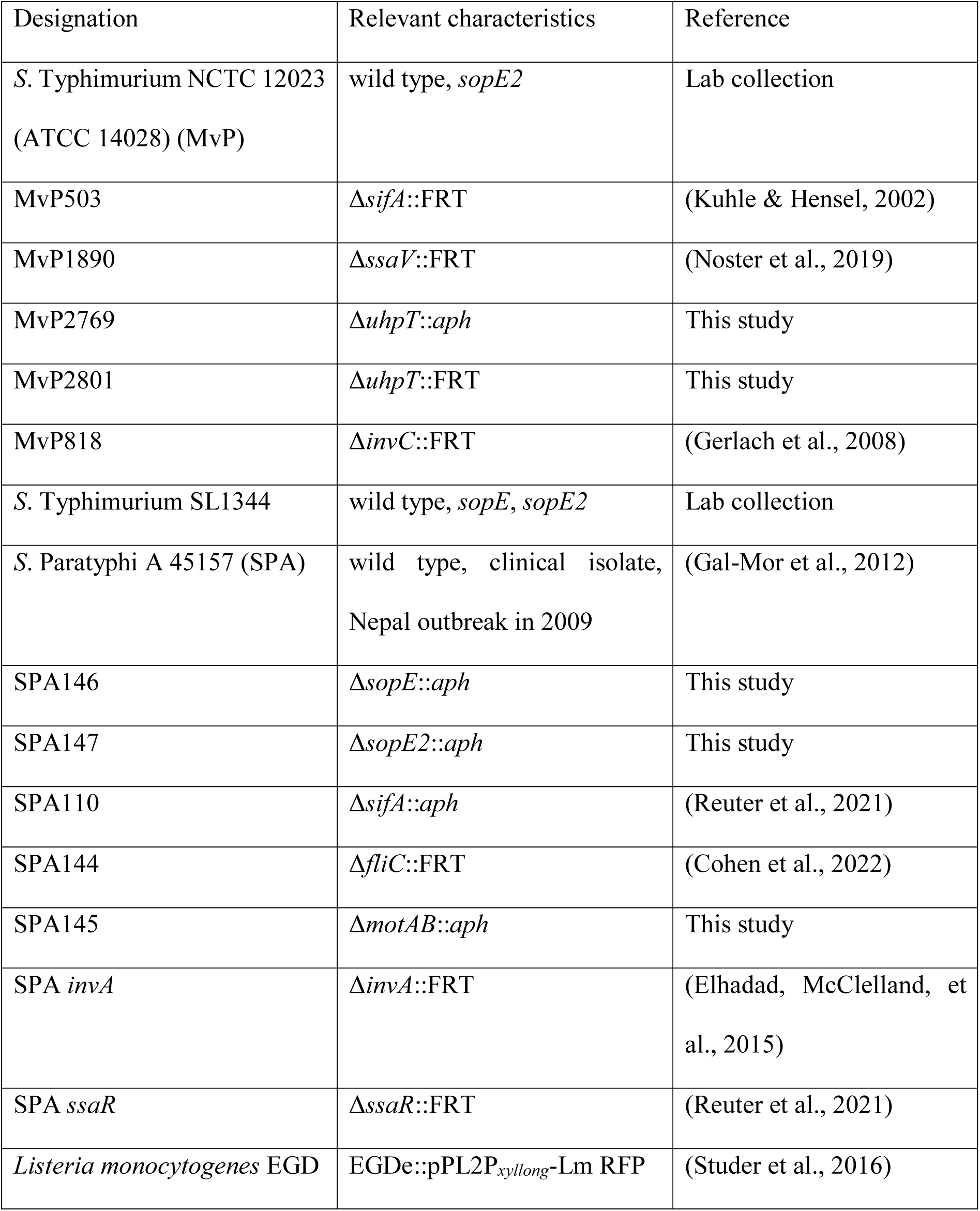
Bacterial strains used in this study.

**Table 2.**
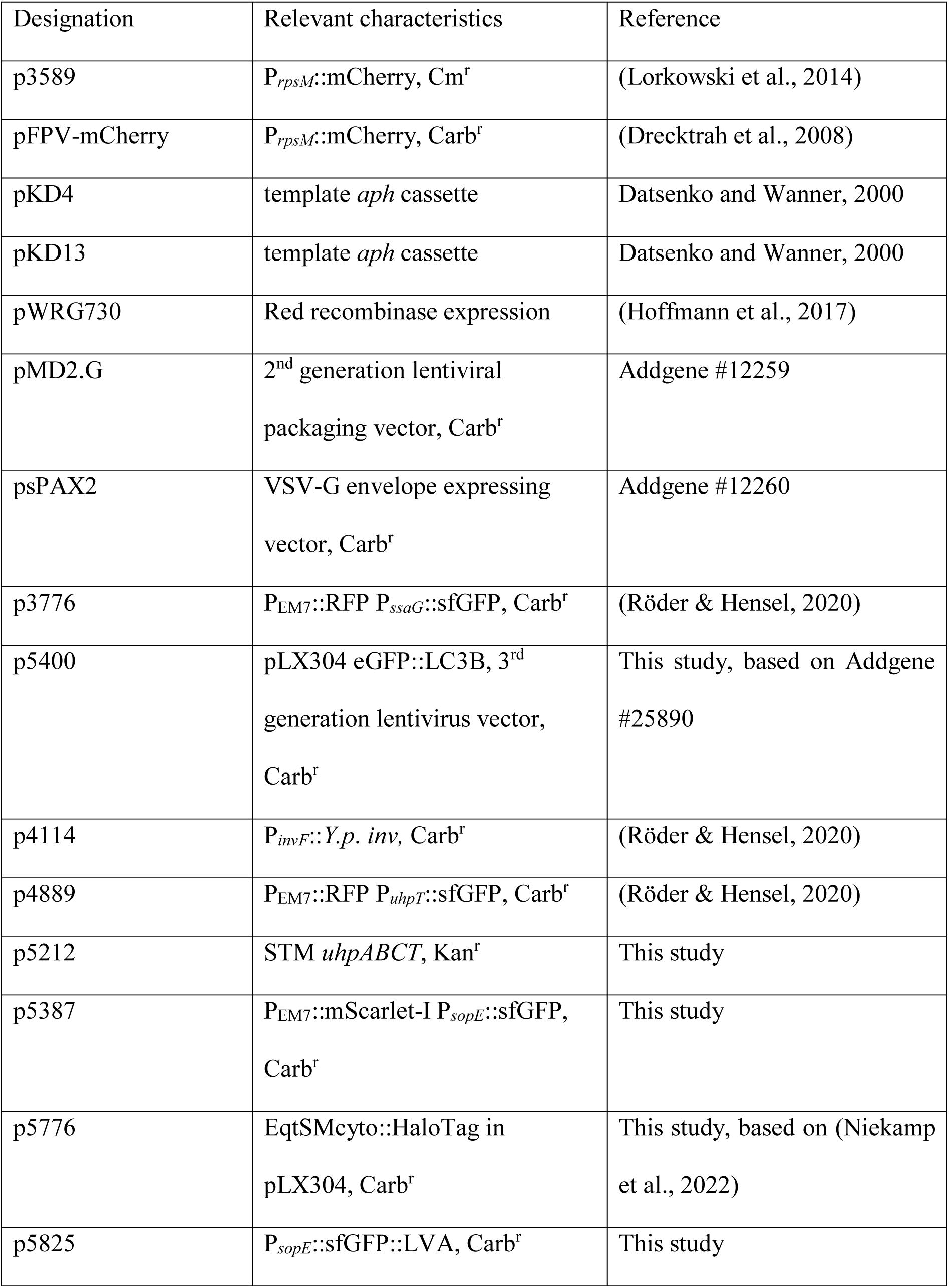
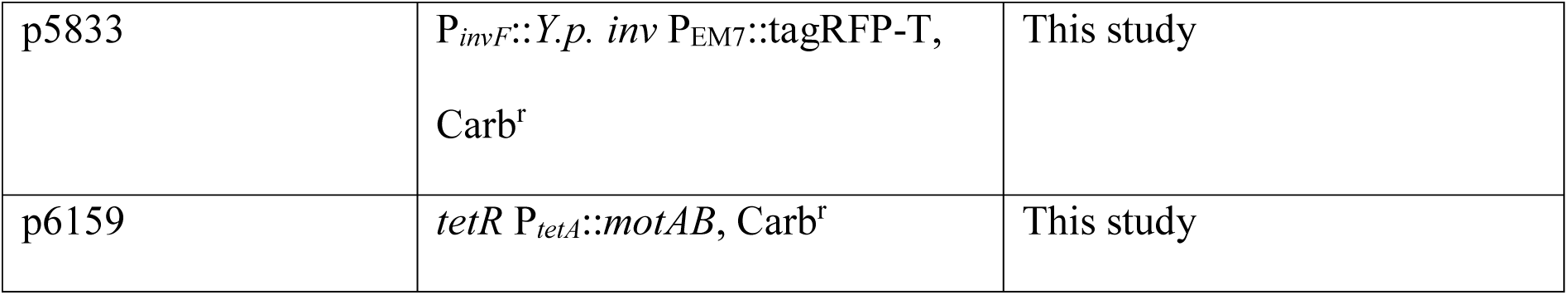
Plasmids used in this study.

| Designation | Relevant characteristics | Reference |
| --- | --- | --- |
| p3589 | P<sub>rpsM</sub>::mCherry, Cm<sup>r</sup> | (Lorkowski et al., 2014) |
| pFPV-mCherry | P<sub>rpsM</sub>::mCherry, Carb<sup>r</sup> | (Drecktrah et al., 2008) |
| pKD4 | template *aph* cassette | Datsenko and Wanner, 2000 |
| pKD13 | template *aph* cassette | Datsenko and Wanner, 2000 |
| pWRG730 | Red recombinase expression | (Hoffmann et al., 2017) |
| pMD2.G | 2<sup>nd</sup> generation lentiviral packaging vector, Carb<sup>r</sup> | Addgene #12259 |
| psPAX2 | VSV-G envelope expressing vector, Carb<sup>r</sup> | Addgene #12260 |
| p3776 | P<sub>EM7</sub>::RFP P<sub>ssaG</sub>::sfGFP, Carb<sup>r</sup> | (Röder & Hensel, 2020) |
| p5400 | pLX304 eGFP::LC3B, 3<sup>rd</sup> generation lentivirus vector, Carb<sup>r</sup> | This study, based on Addgene #25890 |
| p4114 | P<sub>invF</sub>::*Y.p. inv*, Carb<sup>r</sup> | (Röder & Hensel, 2020) |
| p4889 | P<sub>EM7</sub>::RFP P<sub>uhpT</sub>::sfGFP, Carb<sup>r</sup> | (Röder & Hensel, 2020) |
| p5212 | STM *uhpABCT*, Kan<sup>r</sup> | This study |
| p5387 | P<sub>EM7</sub>::mScarlet-I P<sub>sopE</sub>::sfGFP, Carb<sup>r</sup> | This study |
| p5776 | EqtSMcyto::HaloTag in pLX304, Carb<sup>r</sup> | This study, based on (Niekamp et al., 2022) |
| p5825 | P<sub>sopE</sub>::sfGFP::LVA, Carb<sup>r</sup> | This study |
| p5833 | $P_{invF}::Y.p. inv$ $P_{EM7}::tagRFP-T$,<br>Carb<sup>r</sup> | This study |
| p6159 | *tetR* $P_{tetA}::motAB$, Carb<sup>r</sup> | This study |

### Generation of bacterial strains

Isogenic mutant strains were generated by λ Red recombineering for insertion of kanamycin resistance (*aph*) cassettes amplified from template plasmids pKD4 or pKD13 as described before (Chakravortty et al., 2002; Datsenko & Wanner, 2000) using oligonucleotides listed in **Table S1**. Insertion of *aph* cassettes was confirmed by colony PCR. If required, the *aph* cassette was removed by introduction of pE-FLP for FLP-mediated recombination.

### Construction of plasmids

Plasmids were generated by Gibson assembly as described before (Röder et al., 2021; Röder & Hensel, 2020; Schulte et al., 2019) using oligonucleotides listed in **Table S1**. Plasmids used in this study are listed in **Table 2** and were introduced into the respective strains by electroporation.

### Host cell culture and infection

Host cells were cultured and infected as described previously (Cohen et al., 2021). In short, HeLa cells (ATCC no. CCL-2) were seeded in surface-treated 8-well chambered coverslips (ibidi) 24 or 48 h prior to infection to reach ∼80% confluency (∼80,000 cells) and were then used for infection. Cells were continuously maintained in high glucose (4.5 g x l^-1^) DMEM (Merck) supplemented with 10% inactivated fetal calf serum (iFCS, Sigma) at 37 °C in a humidified atmosphere with 5% CO_2_. If indicated, cells were pulse-chased with 100 µg x ml^-1^ fluid phase marker Dextran AlexaFluor 647 (Thermo Fisher Scientific) 16 h prior infection. STM strains for infection experiments were subcultured from an overnight culture (1:31) in fresh LB medium and grown for 3.5 h at 37 °C under aerobic conditions using a roller drum at 60 rpm. SPA strains for infection were grown for 8 h under aerobic conditions as described above and subcultured (1:100) in fresh LB medium and stationary phase subcultures were grown for 16 h under microaerophilic conditions as described in (Elhadad, Desai, et al., 2015). For infection with *Listeria monocytogenes*, overnight cultures grown under aerobic conditions were used. Bacteria were adjusted to an optical density of 0.2 at 600 nm in PBS and used for infection with the respective MOI (between 5 and 90, dependent on strain and experiment). Bacteria were centrifuged onto the cells for 5 min at 500 x *g* to synchronize infection and incubated for 25 min. After washing thrice with PBS, cells were incubated in medium containing 100 µg x ml^-1^ of gentamicin to kill extracellular bacteria. Afterwards, cells were maintained in medium containing 10 µg x ml^-1^ gentamicin until fixation or LCI. To image invasion of *Salmonella* into HeLa cells, cells were infected directly on microscope stage and imaged for ∼1 h.

### Generation of HeLa cells stably expressing LC3B::GFP

HeLa cells stably expressing LC3B-GFP were generated using 3^rd^ generation lentiviral vectors and subsequent fluorescence-activated cell sorting (FACS) (Broad, 2015; Institute, 2015; TRC, 2015a, 2015b). Supernatant containing lentivirus was generated by seeding 3.8 x 10^5^ cells x ml^-1^ HEK 293FT cells (Invitrogen R700-07) in 10 cm tissue culture plates (10 ml per plate, TPP) in antibiotic-free growth media (high glucose DMEM + 10% iFCS). After 24 h, cells should be 70-80% confluent and were then transfected with lentiviral packaging plasmid (9 µg psPAX2), envelope plasmid (0.9 µg pMD2.G) and pLX304 carrying eGFP-LC3B (9 µg) using Opti-MEM (Fisher Scientific) and FuGENE HD transfection reagent (Promega). Transfection mix was incubated for 30 min at room temperature before added to cells dropwise. Cells were incubated for at least 18 h and medium was changed to 15 ml growth medium containing 30% iFCS. Medium containing lentivirus was harvested after 24 h and 48 h and centrifuged at 350 x *g* for 5 min to pellet any residual packaging cells. Supernatant was stored in sterile polypropylene storage tubes and stored at -80 °C. Half of the harvested supernatant was filtered using fast flow & low binding filters (Merck Millipore).

HeLa cells were seeded in 24-well plates (1 x 10^5^ cells x ml^-1^) and incubated for 24 h before 8 µg x ml^-1^ Polybrene (Sigma) was added to the cells. Filtered supernatant containing lentivirus was added to cells 48 h after seeding. 48 h after lentivirus was added to cells, medium was exchanged to DMEM containing 10 µg x ml^-1^ Blasticidin (Invitrogen) and incubated for 72 h. Medium was changed to DMEM without antibiotics to allow growth for 24 h before cells were detached for culture in tissue culture flasks (TPP) to obtain sufficient cells numbers for FACS.

For FACS of lentivirus transfected HeLa cells, cells were detached from tissue culture flask and resuspended in appropriate amount of medium to reach concentration of 1 x 10^7^ cells x ml^-1^. A 100 µm cell strainer (BD Falcon) was used to prevent aggregation of cells prior to FACS. Cells were sorted for GFP fluorescence with BD FACS Aria. A mixed population of cells with different GFP expression levels was obtained. GFP-positive cells were cultured again in tissue culture flasks before a second round of FACS with Sony SH800S was conducted. Single cells were sorted into 96-well plates to obtain single clones of LC3B-GFP expressing cells. Cell clones were analyzed regarding GFP intensity, cell division, and phenotypes of intracellular *Salmonella* prior to use in further experiments.

### Live cell imaging (LCI)

Prior to LCI, medium of infected HeLa cells was changed to high glucose DMEM without phenol red supplemented with 30 mM HEPES. LCI was performed at 37 °C and an atmosphere of 5% CO_2_ with Cell Observer microscope (Zeiss) equipped with Yokogawa Spinning Disc Unit CSU X1a5000, an incubation chamber, 63x objective (α-Plan-Apochromat, NA 1.46) and 40x objective (Plan-Apochromat, NA 1.4), two ORCA Flash 4.0 V3 cameras (Hamamatsu) and appropriate filters for the respective fluorescence proteins or dyes.

### Correlative light and electron microscopy (CLEM)

CLEM of HeLa LC3B-GFP infected by SPA was performed as previously described (Krieger et al., 2014). Briefly, HeLa LC3B-GFP cells were grown on MatTek dishes with gridded coverslips. Cells were fixed with 4% paraformaldehyde (Electron Microscopy Sciences) and 0.1% glutaraldehyde (Electron Microscopy Sciences) in 200 mM HEPES directly on microscope stage during imaging of cytosolic motile SPA. Cells were incubated for 30 min and were then rinsed thrice with 200 mM HEPES buffer. Cells were subsequently fixed with 2.5% glutaraldehyde in 200 mM HEPES for 1 h, washed thrice with buffer and post-fixed with 2% osmium-tetroxide (Electron Microscopy Sciences), 0.1% ruthenium red (Applichem) and 1.5% potassium ferrocyanide (Sigma) in 200 mM HEPES on ice. After several washing steps, the cells were dehydrated in a cold graded series of ethanol and finally one rinse in anhydrous ethanol and two rinses in anhydrous acetone at room temperature. Infiltration was performed with increasing concentrations of EPON812 (Sigma) in anhydrous acetone. Utilizing the coordinate system, the ROI from light microscopy was relocated on the EPON block, trimmed and 70 nm serial sections were cut with an ultramicrotome (Leica EM UC7, Leica, Wetzlar, Germany) and collected on formvar-coated copper slot grids (Plano). Grids were post-stained for 30 min with 2% uranyl acetate (Roth) and 20 min with 3% lead citrate (Leica). TEM images were acquired using a Zeiss Leo 912 Omega (Zeiss, Oberkochen, Germany) equipped with a CCD camera (TRS, Moorenwies, Germany). Overlays of the light and electron microscopic images were generated using LAS AF (Leica) and Photoshop CS6 (Adobe).

### Tracking analyses

For microscopic tracking analyses, bacterial cultures were imaged with Cell Observer microscope (Zeiss) equipped with 100x objective (α-Plan-Apochromat, NA 1.46) and CoolSNAP camera at about 8 frames per second (fps) over 1 min. For tracking of intracellular SPA, infected cells were imaged with Cell Observer microscope (Zeiss) equipped with Yokogawa Spinning Disc Unit CSU X1a5000, an incubation chamber, 63x objective (α-Plan-Apochromat, NA 1.46), two ORCA Flash 4.0 V3 cameras (Hamamatsu) and appropriate filters for the respective fluorescence proteins at about 18 fps over 1 min. Tracking analyses were performed in Fiji with TrackMate v5.0.2 plugin (Tinevez et al., 2017) with minimal displacement threshold of 1 µm. For tracking of intracellular *L.m.*, infected cells were imaged with setup as described for intracellular SPA at ∼0.1 fps over 10-30 min. Tracking analyses were performed in FIJI with manual tracking plugin.

### Analyses of LC3B decoration

For analyses of LC3B-decoration of intracellular SPA, randomly selected infected cells were imaged 3-4 h p.i. as described above and were analyzed manually regarding LC3B-GFP signal at SPA cell bodies. 100 µg x ml^-1^ cefotaxime and ciprofloxacin were added 2 h p.i. if indicated.

For correlation of maximum velocity and LC3B decoration of cytosolic SPA, individual bacteria were analyzed regarding LC3B-GFP fluorescence intensity using ZEISS Efficient Navigation (ZEN) software. Manual tracking with the same bacteria was performed as described above.

### Transfection of HeLa cells with Halo-tagged EqtSM

HeLa cells were maintained and infected as described above. In short, cells were seeded in 8-wells 2 d prior infection. Cells were transfected with Halo-tagged EqtSM expression construct 1 d prior infection using FuGENE HD in 1:2 ratio. Infected cells were labeled with 100 nM Janelia Fluor 646 HaloTag ligand (Promega, GA1120) for 30 min, washed five times with PBS and subsequently imaged using the Zeiss Cell Observer SD microscope set-up.

## Suppl. Materials and Methods

### Immuno-staining and fluorescence microscopy

For immuno-staining of HeLa LAMP1-GFP cells infected by SPA, cells were seeded in 24-well plates (TPP) on coverslips 24 h or 48 h prior infection to reach ∼80% confluency (∼180,000 cells). Cultivation of bacteria and infection of HeLa cells was carried out as described above. Infected cells were fixed with 3% PFA at the desired time point. After washing thrice with PBS, cells were incubated in blocking solution (2% goat serum, 2% BSA, 0.1% saponin in PBS) for 30 min. Next, cells were incubated for 1 h at RT with mouse monoclonal antibodies against Ubiquitin (Biomol PW 8810). After washing cells thrice with PBS, cells were incubated with primary antibodies against *Salmonella* O-Antigen of SPA (BD Difco 229471) for 1 h. After washing thrice with PBS, cells were stained with the appropriate secondary antibodies for 1 h (Invitrogen A11031, Dianova 111-607-003). After three washing steps with PBS, coverslips were mounted with Fluoroshield (Sigma) and sealed with Entellan (Merck). Microscopy of fixed samples was performed with Leica SP5 confocal laser-scanning microscope using 100x objective (HCX PL APO CS, NA 1.4-0.7) and polychroic mirror TD 488/543/633.

### Flow cytometry

The assay was performed as described previously (Reuter et al., 2021). Briefly, HeLa cells were seeded in 12-well plates (TPP) 48 h prior infection to reach confluency (∼4 x 10^5^ cells per well) on the day of infection. Cells were infected with STM and SPA cultures at MOI 30 as described above. The bacterial strains harbored fluorescence reporter plasmids. At 1 h and 3 h p.i., cells were washed thrice with PBS, detached from culture plates, fixed with 3% PFA and analyzed by flow cytometry using Attune NxT cytometer. At least 3,000 infected cells were measured and analyzed regarding expression of fluorescence reporter.

### Gentamicin protection assays

The assay was performed as described before (Kuhle & Hensel, 2002). Briefly, HeLa cells were seeded in 24-well plates (TPP) 48 h prior infection to reach confluency (∼2 x 10^5^ cells per well) on the day of infection. Cells were infected with STM and SPA cultures at MOI 5 as described above. Cells were washed thrice with PBS and lysed using 0.1% Triton X-100 at 1 h and 16 h p.i. Colony forming units were determined by plating serial dilutions of lysates and inoculum on Mueller-Hinton II agar and incubated overnight at 37 °C. The percentage of internalized bacteria of the inoculum (from subcultures) and the replication rate of intracellular bacteria was calculated.

## Suppl. Tables

**Table S1.**
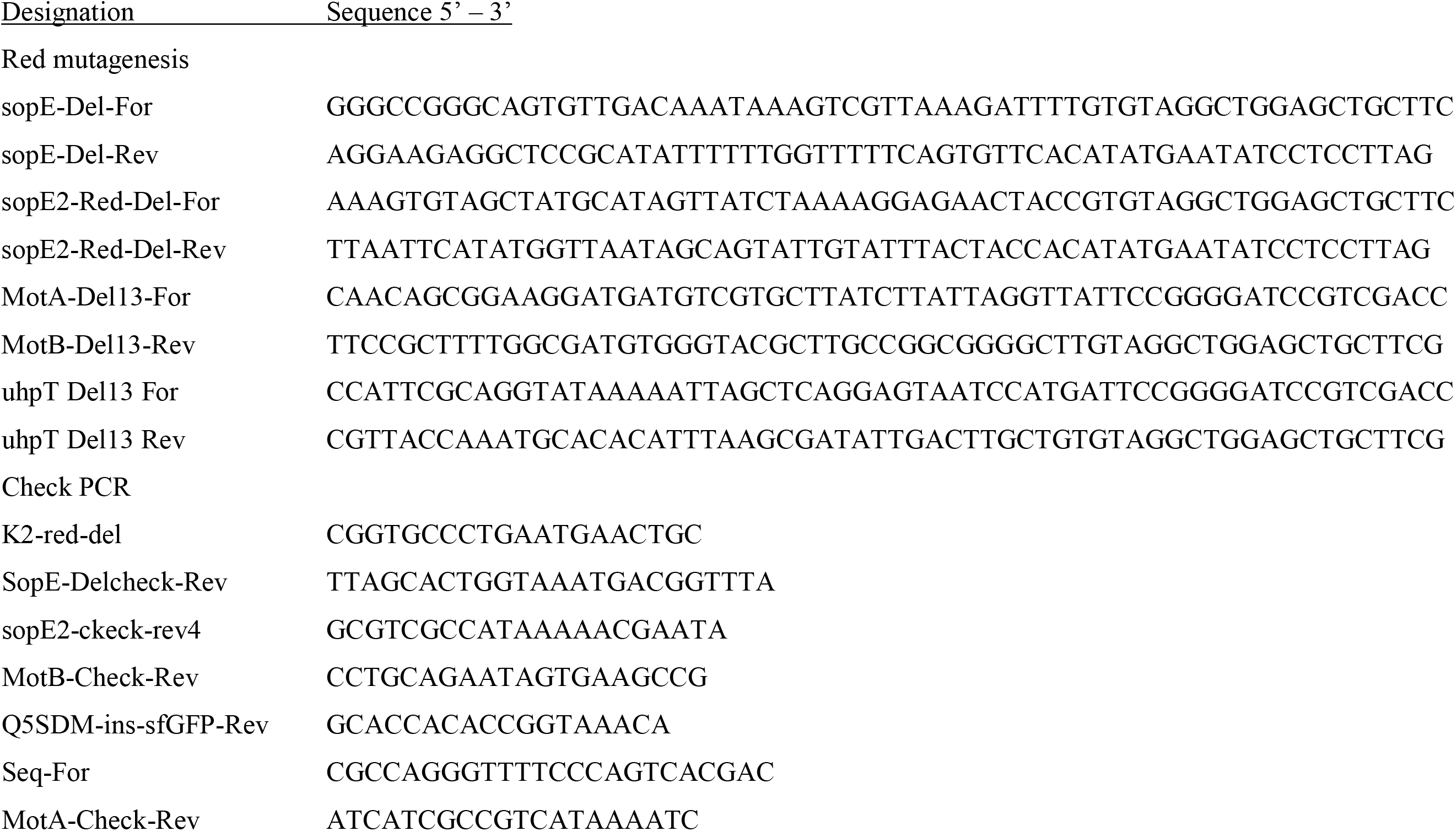
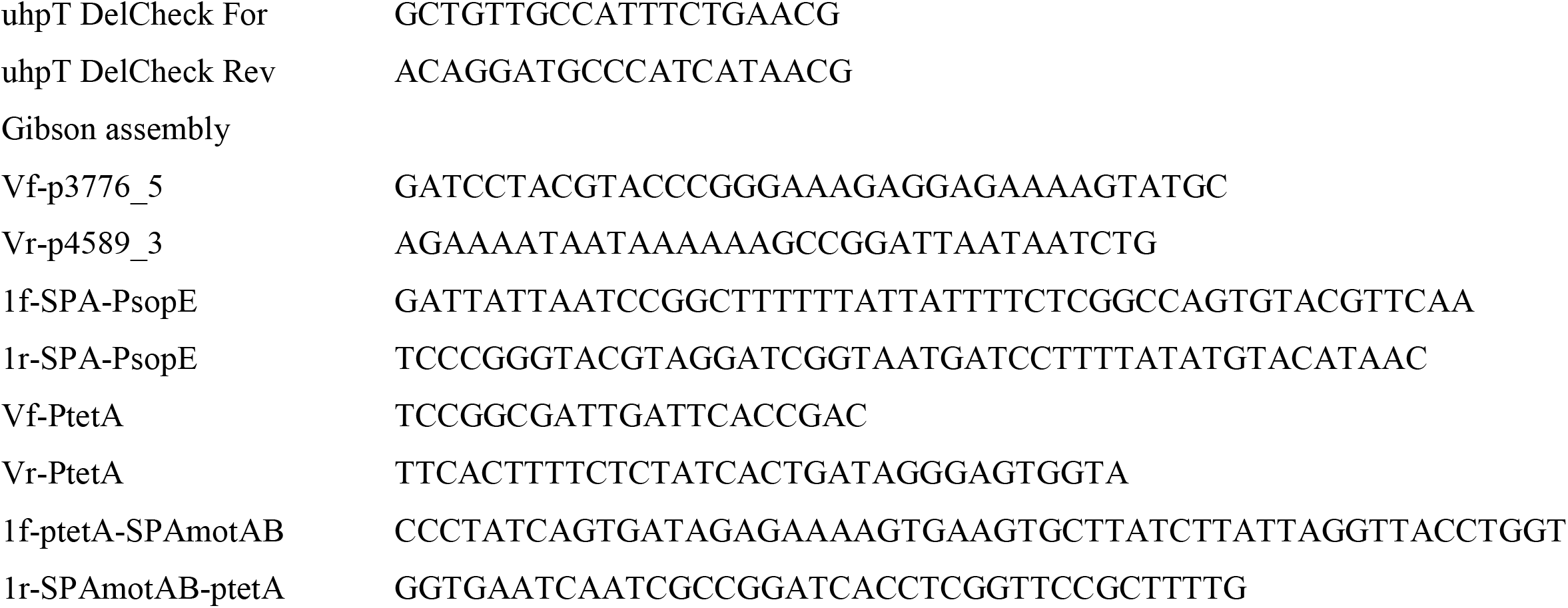
Oligonucleotides used in this study.

**Fig. S1.**
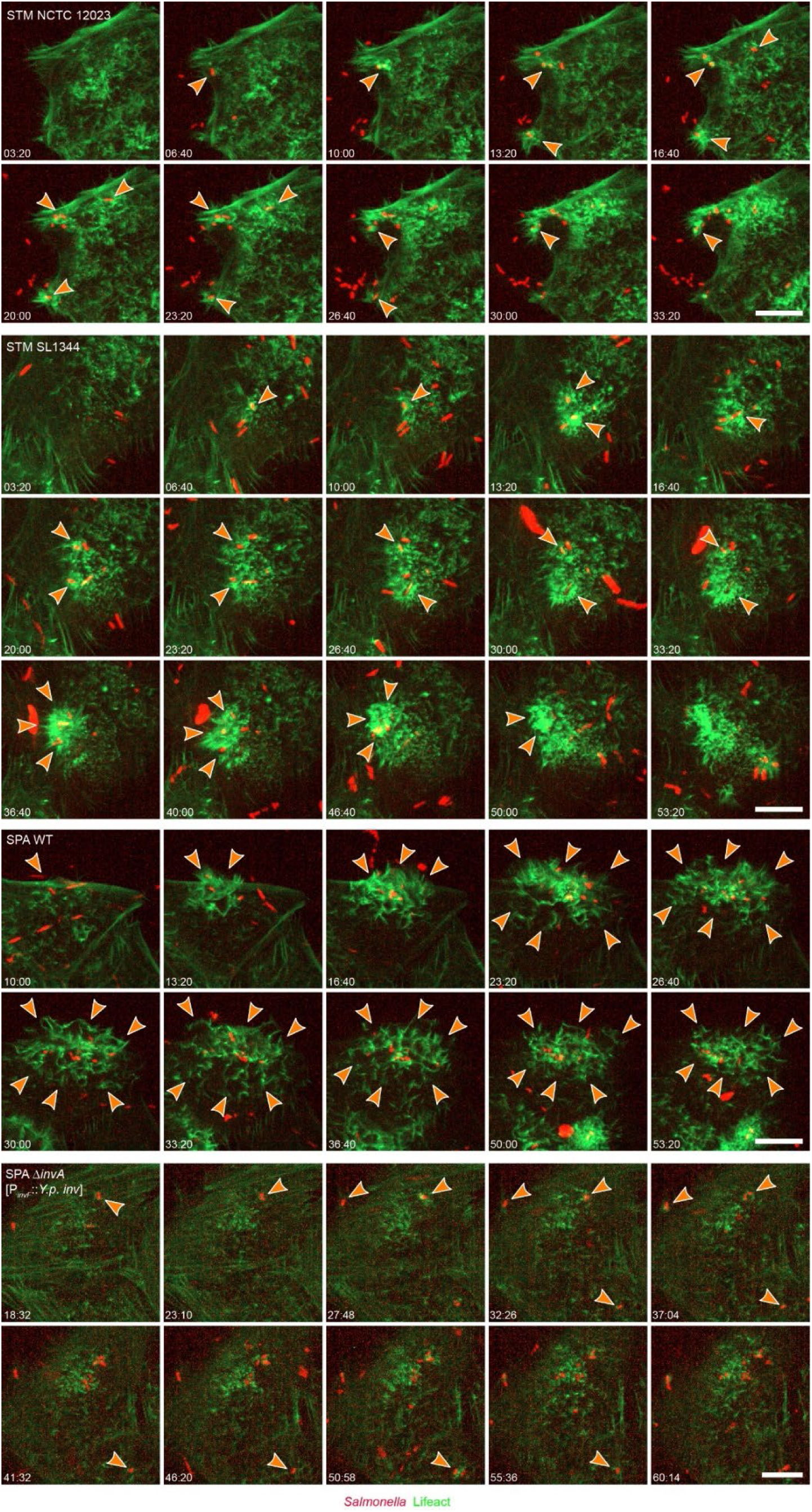
SPA trigger invasion induces massive membrane ruffles and mediates highly cooperative invasion. HeLa cells stably expressing Lifeact-eGFP (green) were infected as indicated with STM NCTC 12023, STM SL1344, SPA WT, or SPA Δ*invA* [P_*invF*_::*Y.p. inv*] expressing mCherry (red) at MOI 75. Cells were infected on microscope stage and imaged for 1 h by spinning disc confocal microscopy. Still images from representative time-lapse series are shown, corresponding with the quantitative analyses shown in Fig. 2. Time stamp indicating min:sec. Sites of ongoing invasion used for quantification of number of bacteria are indicated by arrowheads. Scale bars: 10 µm.

**Fig. S2.**
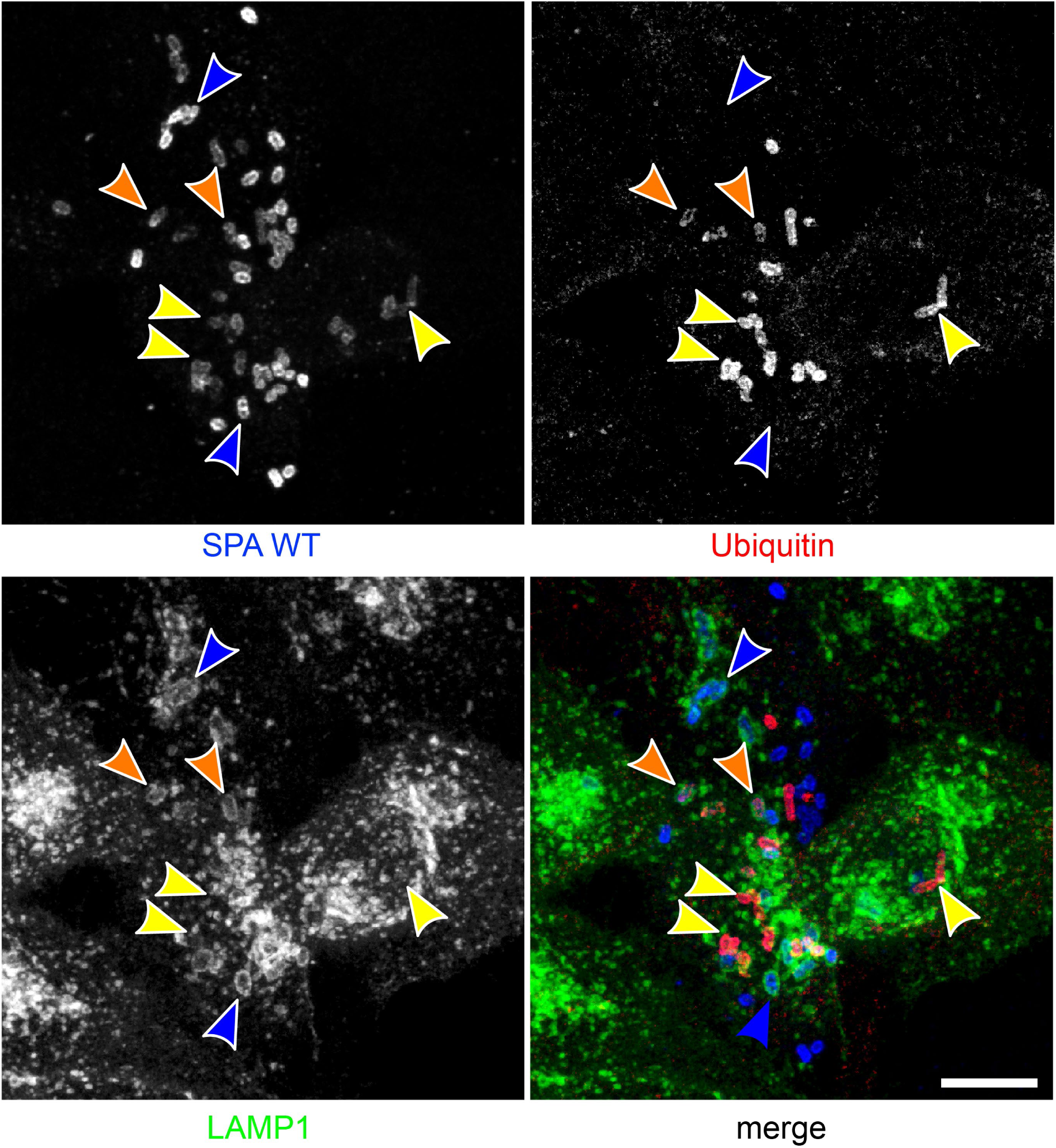
Ubiquitylation of cytosolic SPA. HeLa cells expressing LAMP1-GFP (green) were infected with SPA WT at MOI 30. Cells were fixed at 7 h p.i. and immunolabeled for SPA LPS (blue) and ubiquitin (red) prior to confocal microscopy. A representative maximum intensity projection of an infected host cell is shown. Arrowheads indicate SPA in SCV without ubiquitin signal (white), cytosolic SPA with ubiquitin signal (yellow), and SPA within partially intact SCV and ubiquitin signal (orange), Scale bar: 10 µm.

**Fig. S3.**
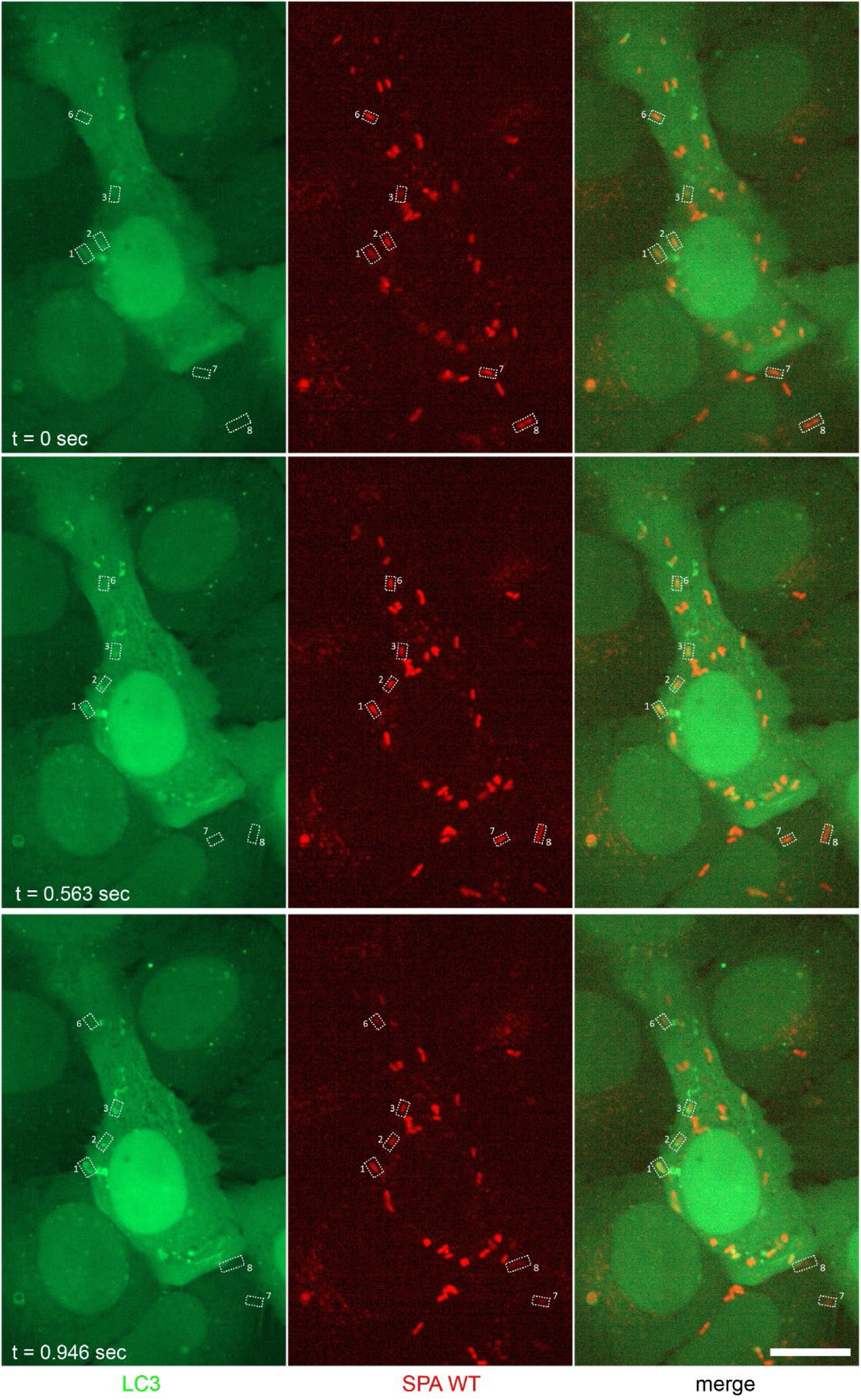
Single cell quantification of LC3B-GFP fluorescence intensity and velocity of cytosolic SPA. HeLa LC3B-GFP cells were infected with SPA WT expressing mCherry at MOI 30. At 3-4 h p.i. cells were imaged and cytosolic SPA were identified (boxes). For single SPA cells LC3B-GFP fluorescence signal intensity was recorded, and velocity was determined by single cell tracking. Three time points of a representative time lapse series are shown, and quantitative data are displayed in Fig. 6. Images shown are single Z-planes from a stack. Scale bar: 10 µm.

**Fig. S4.**
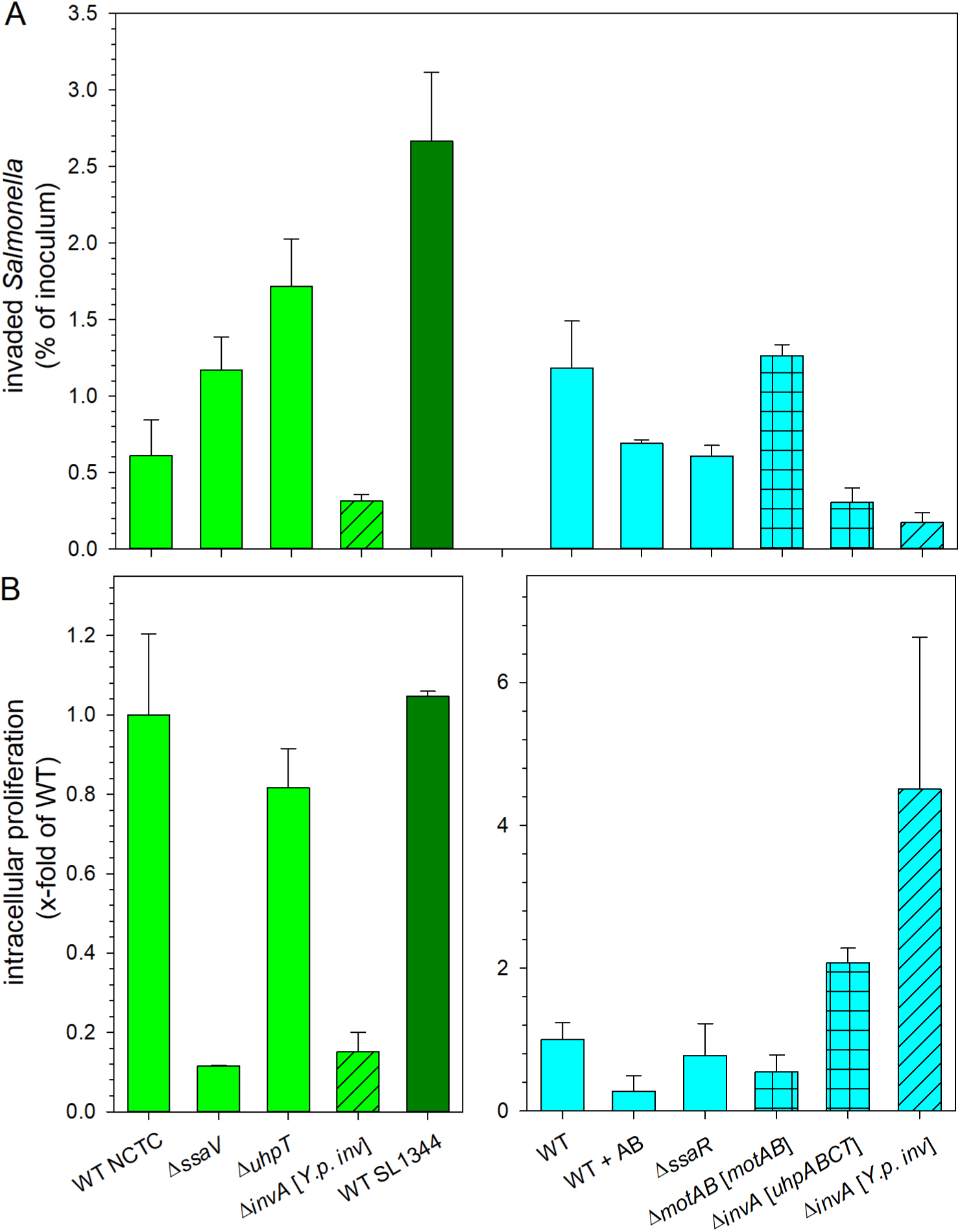
Host cell invasion and intracellular replication of various STM and SPA strains. HeLa cells were infected with STM (green) or SPA (cyan) WT or isogenic mutant strains at MOI 5. Presence of plasmids for expression *Y. pseudotuberculosis inv*, P_*tetA*_::*motAB*, or STM *uhpABCT* is indicated by hatched bars. AB indicates addition of antibiotics cefotaxime and ciprofloxacin. Invasion (A) and intracellular replication (B) were determined by gentamicin protection assays comparing CFU counts of inoculum with lysates 1 h and 16 h p.i. Depicted are means and standard deviations of triplicate samples from one biological replicate performed.

## Suppl. Movie Captions

### Movie 1: Live cell imaging of STM NCTC 12023 trigger invasion

HeLa cells stably expressing Lifeact-GFP (green) were infected with STM NCTC 12023 WT expressing mCherry (red) and imaged over 1 h. The movie successively shows two individual clips. Time stamps in upper corner indicates h:min:sec. Scale bar: 10 µm.

Link to movie: https://myshare.uni-osnabrueck.de/f/01372357bfbb44c49087/

### Movie 2: Live cell imaging of STM SL1344 trigger invasion

HeLa cells stably expressing Lifeact-GFP (green) were infected with STM SL1344 WT expressing mCherry (red) and imaged over 1 h. The movie successively shows three individual clips. Time stamps in upper corner indicates h:min:sec. Scale bar: 10 µm.

Link to movie: https://myshare.uni-osnabrueck.de/f/7c96d08641e24bc3b293/

### Movie 3: Live cell imaging of SPA WT trigger invasion

HeLa cells stably expressing Lifeact-GFP (green) were infected with SPA WT expressing mCherry (red) and imaged over 1 h. The movie successively shows two individual clips. Time stamps in upper corner indicates h:min:sec. Scale bar: 10 µm.

Link to movie: https://myshare.uni-osnabrueck.de/f/62a3a98281dc496a9b04/

### Movie 4: Live cell imaging of SPA zipper invasion

HeLa cells stably expressing Lifeact-GFP (green) were infected with SPA *invA* [*Y.p. inv*] expressing mCherry (red) and imaged over 1 h. The movie successively shows three individual clips. Time stamps in upper corner indicates h:min:sec. Scale bar: 10 µm.

Link to movie: https://myshare.uni-osnabrueck.de/f/5ae1904709b64a679b56/

### Movie 5: Live cell imaging of intracellular SPA WT targeted by LC3B

HeLa cells stably expressing LC3B-GFP (green) were infected with SPA WT expressing mCherry (red) and imaged every two minutes over two hours. The movie successively shows three individual clips with various degrees of xenophagic capture. Time stamps indicates minutes. Scale bar: 10 µm.

Link to movie: https://myshare.uni-osnabrueck.de/f/bbd7f3927781420382dc/

### Movie 6: Real-time imaging of intracellular SPA WT

HeLa cells stably expressing LC3B-GFP (green) were infected with SPA WT expressing mCherry (red) and pulse-chased with Dextran AlexaFluor 647 (blue) if indicated, and imaged at frame rates of 2-3 frames per second. Movie successively shows thee individual clips with various degrees of xenophagic capture. Time stamps indicates seconds. Scale bar: 10 µm.

Link to movie: https://myshare.uni-osnabrueck.de/f/50d491e54e6a48c68c5a/

## Notes

https://myshare.uni-osnabrueck.de/f/01372357bfbb44c49087/

https://myshare.uni-osnabrueck.de/f/7c96d08641e24bc3b293/

https://myshare.uni-osnabrueck.de/f/62a3a98281dc496a9b04/

https://myshare.uni-osnabrueck.de/f/5ae1904709b64a679b56/

https://myshare.uni-osnabrueck.de/f/bbd7f3927781420382dc/

https://myshare.uni-osnabrueck.de/f/50d491e54e6a48c68c5a/

### Competing Interest Statement

The authors have declared no competing interest.

### Summary of Updates

Figure 1 was replaced by a new version that contains a legend.

## References

Birmingham, C. L., Canadien, V., Gouin, E., Troy, E. B., Yoshimori, T., Cossart, P., Higgins, D. E., & Brumell, J. H. (2007). *Listeria monocytogenes* evades killing by autophagy during colonization of host cells. Autophagy, 3(5), 442–451. https://doi.org/10.4161/auto.4450

Broad, I. (2015). Lentivirus production of ShRNA, CRISPR, or ORF-pLX clones in 10 cm dishes or 6-well plates [TRC Laboratory Protocols]. https://portals.broadinstitute.org/gpp/public/resources/protocols

Brumell, J. H., Tang, P., Zaharik, M. L., & Finlay, B. B. (2002). Disruption of the *Salmonella*-containing vacuole leads to increased replication of *Salmonella enterica* serovar typhimurium in the cytosol of epithelial cells. Infect Immun, 70(6), 3264–3270. https://www.ncbi.nlm.nih.gov/pubmed/12011022

Castanheira, S., & Garcia-Del Portillo, F. (2017). *Salmonella* Populations inside Host Cells. Front Cell Infect Microbiol, 7, 432. https://doi.org/10.3389/fcimb.2017.00432

Chakravortty, D., Hansen-Wester, I., & Hensel, M. (2002). *Salmonella* pathogenicity island 2 mediates protection of intracellular *Salmonella* from reactive nitrogen intermediates. J Exp Med, 195(9), 1155–1166. https://doi.org/10.1084/jem.20011547

Cohen, E., Azriel, S., Auster, O., Gal, A., Zitronblat, C., Mikhlin, S., Scharte, F., Hensel, M., Rahav, G., & Gal-Mor, O. (2021). Pathoadaptation of the passerine-associated *Salmonella enterica* serovar Typhimurium lineage to the avian host. PLoS Pathog, 17(3), e1009451. https://doi.org/10.1371/journal.ppat.1009451

Cohen, H., Hoede, C., Scharte, F., Coluzzi, C., Cohen, E., Shomer, I., Mallet, L., Holbert, S., Serre, R. F., Schiex, T., Virlogeux-Payant, I., Grassl, G. A., Hensel, M., Chiapello, H., & Gal-Mor, O. (2022). Intracellular Salmonella Paratyphi A is motile and differs in the expression of flagella-chemotaxis, SPI-1 and carbon utilization pathways in comparison to intracellular S. Typhimurium. PLoS Pathog, 18(4), e1010425. https://doi.org/10.1371/journal.ppat.1010425

Connor, M. G., Pulsifer, A. R., Chung, D., Rouchka, E. C., Ceresa, B. K., & Lawrenz, M. B. (2018). *Yersinia pestis* Targets the Host Endosome Recycling Pathway during the Biogenesis of the *Yersinia*-Containing Vacuole To Avoid Killing by Macrophages. mBio, 9(1). https://doi.org/10.1128/mBio.01800-17

Datsenko, K. A., & Wanner, B. L. (2000). One-step inactivation of chromosomal genes in *Escherichia coli* K-12 using PCR products. Proc Natl Acad Sci U S A, 97(12), 6640–6645. https://doi.org/10.1073/pnas.120163297

Deng, Y., Rivera-Molina, F. E., Toomre, D. K., & Burd, C. G. (2016). Sphingomyelin is sorted at the trans Golgi network into a distinct class of secretory vesicle. Proc Natl Acad Sci U S A, 113(24), 6677–6682. https://doi.org/10.1073/pnas.1602875113

Dougan, G., & Baker, S. (2014). *Salmonella enterica* serovar Typhi and the pathogenesis of typhoid fever. Annu Rev Microbiol, 68, 317–336. https://doi.org/10.1146/annurev-micro-091313-103739

Dowd, G. C., Mortuza, R., & Ireton, K. (2021). Molecular Mechanisms of Intercellular Dissemination of Bacterial Pathogens. Trends Microbiol, 29(2), 127–141. https://doi.org/10.1016/j.tim.2020.06.008

Drecktrah, D., Levine-Wilkinson, S., Dam, T., Winfree, S., Knodler, L. A., Schroer, T. A., & Steele-Mortimer, O. (2008). Dynamic behavior of *Salmonella*-induced membrane tubules in epithelial cells. Traffic, 9(12), 2117–2129. https://doi.org/10.1111/j.1600-0854.2008.00830.x

Elhadad, D., Desai, P., Rahav, G., McClelland, M., & Gal-Mor, O. (2015). Flagellin Is Required for Host Cell Invasion and Normal *Salmonella* Pathogenicity Island 1 Expression by *Salmonella enterica* Serovar Paratyphi A. Infect Immun, 83(9), 3355–3368. https://doi.org/10.1128/IAI.00468-15

Elhadad, D., McClelland, M., Rahav, G., & Gal-Mor, O. (2015). Feverlike Temperature is a Virulence Regulatory Cue Controlling the Motility and Host Cell Entry of Typhoidal *Salmonella*. J Infect Dis, 212(1), 147–156. https://doi.org/10.1093/infdis/jiu663

Ellison, C. J., Kukulski, W., Boyle, K. B., Munro, S., & Randow, F. (2020). Transbilayer Movement of Sphingomyelin Precedes Catastrophic Breakage of Enterobacteria-Containing Vacuoles. Curr Biol, 30(15), 2974–2983 e2976. https://doi.org/10.1016/j.cub.2020.05.083

Forest, C. G., Ferraro, E., Sabbagh, S. C., & Daigle, F. (2010). Intracellular survival of *Salmonella enterica* serovar Typhi in human macrophages is independent of *Salmonella* pathogenicity island (SPI)-2. Microbiology, 156(Pt 12), 3689–3698. https://doi.org/10.1099/mic.0.041624-0

Gal-Mor, O., Suez, J., Elhadad, D., Porwollik, S., Leshem, E., Valinsky, L., McClelland, M., Schwartz, E., & Rahav, G. (2012). Molecular and cellular characterization of a *Salmonella enterica* serovar Paratyphi a outbreak strain and the human immune response to infection. Clin Vaccine Immunol, 19(2), 146–156. https://doi.org/10.1128/CVI.05468-11

Gerlach, R. G., Claudio, N., Rohde, M., Jäckel, D., Wagner, C., & Hensel, M. (2008). Cooperation of *Salmonella* pathogenicity islands 1 and 4 is required to breach epithelial barriers. Cell Microbiol, 10(11), 2364–2376. https://doi.org/10.1111/j.1462-5822.2008.01218.x

Göser, V., Kehl, A., Röder, J., & Hensel, M. (2020). Role of the ESCRT-III complex in controlling integrity of the *Salmonella*-containing vacuole. Cell Microbiol, 22(6), e13176. https://doi.org/10.1111/cmi.13176

Hiyoshi, H., Tiffany, C. R., Bronner, D. N., & Baumler, A. J. (2018). Typhoidal *Salmonella* serovars: ecological opportunity and the evolution of a new pathovar. FEMS Microbiol Rev, 42(4), 527–541. https://doi.org/10.1093/femsre/fuy024

Hiyoshi, H., Wangdi, T., Lock, G., Saechao, C., Raffatellu, M., Cobb, B. A., & Baumler, A. J. (2018). Mechanisms to Evade the Phagocyte Respiratory Burst Arose by Convergent Evolution in Typhoidal Salmonella Serovars. Cell Rep, 22(7), 1787–1797. https://doi.org/10.1016/j.celrep.2018.01.016

Hoffmann, S., Schmidt, C., Walter, S., Bender, J. K., & Gerlach, R. G. (2017). Scarless deletion of up to seven methyl-accepting chemotaxis genes with an optimized method highlights key function of CheM in *Salmonella* Typhimurium. PLoS One, 12(2), e0172630. https://doi.org/10.1371/journal.pone.0172630

Holt, K. E., Thomson, N. R., Wain, J., Langridge, G. C., Hasan, R., Bhutta, Z. A., Quail, M. A., Norbertczak, H., Walker, D., Simmonds, M., White, B., Bason, N., Mungall, K., Dougan, G., & Parkhill, J. (2009). Pseudogene accumulation in the evolutionary histories of *Salmonella enterica* serovars Paratyphi A and Typhi. BMC Genomics, 10, 36. https://doi.org/10.1186/1471-2164-10-36

Institute, T. R. C. T. B. (2015). Lentivirus production of ShRNA, CRISPR, or ORF-pLX clones in 10 cm dishes or 6-well plates [TRC Laboratory Protocols]. https://portals.broadinstitute.org/gpp/public/resources/protocols

Jennings, E., Thurston, T. L. M., & Holden, D. W. (2017). *Salmonella* SPI-2 Type III Secretion System Effectors: Molecular Mechanisms And Physiological Consequences. Cell Host Microbe, 22(2), 217–231. https://doi.org/10.1016/j.chom.2017.07.009

Johnson, R., Byrne, A., Berger, C. N., Klemm, E., Crepin, V. F., Dougan, G., & Frankel, G. (2017). The Type III Secretion System Effector SptP of *Salmonella enterica* Serovar Typhi. J Bacteriol, 199(4). https://doi.org/10.1128/JB.00647-16

Knodler, L. A., Vallance, B. A., Celli, J., Winfree, S., Hansen, B., Montero, M., & Steele-Mortimer, O. (2010). Dissemination of invasive *Salmonella* via bacterial-induced extrusion of mucosal epithelia. Proc Natl Acad Sci U S A, 107(41), 17733–17738. https://doi.org/10.1073/pnas.1006098107

Krieger, V., Liebl, D., Zhang, Y., Rajashekar, R., Chlanda, P., Giesker, K., Chikkaballi, D., & Hensel, M. (2014). Reorganization of the endosomal system in *Salmonella*-infected cells: the ultrastructure of *Salmonella*-induced tubular compartments. PLoS Pathog, 10(9), e1004374. https://doi.org/10.1371/journal.ppat.1004374

Kubori, T., & Galan, J. E. (2003). Temporal regulation of *Salmonella* virulence effector function by proteasome-dependent protein degradation. Cell, 115(3), 333–342. 14636560

Kuhle, V., & Hensel, M. (2002). SseF and SseG are translocated effectors of the type III secretion system of *Salmonella* pathogenicity island 2 that modulate aggregation of endosomal compartments. Cell Microbiol, 4(12), 813–824. https://doi.org/10.1046/j.1462-5822.2002.00234.x

Lacayo, C. I., & Theriot, J. A. (2004). *Listeria monocytogenes* actin-based motility varies depending on subcellular location: a kinematic probe for cytoarchitecture. Mol Biol Cell, 15(5), 2164–2175. https://doi.org/10.1091/mbc.e03-10-0747

Lemarignier, M., & Pizarro-Cerda, J. (2020). Autophagy and Intracellular Membrane Trafficking Subversion by Pathogenic *Yersinia* Species. Biomolecules, 10(12). https://doi.org/10.3390/biom10121637

Liu, W., Zhou, Y., Peng, T., Zhou, P., Ding, X., Li, Z., Zhong, H., Xu, Y., Chen, S., Hang, H. C., & Shao, F. (2018). N(epsilon)-fatty acylation of multiple membrane-associated proteins by *Shigella* IcsB effector to modulate host function. Nat Microbiol, 3(9), 996–1009. https://doi.org/10.1038/s41564-018-0215-6

Lorkowski, M., Felipe-Lopez, A., Danzer, C. A., Hansmeier, N., & Hensel, M. (2014). *Salmonella enterica* invasion of polarized epithelial cells is a highly cooperative effort. Infect Immun, 82(6), 2657–2667. https://doi.org/10.1128/IAI.00023-14

McClelland, M., Sanderson, K. E., Clifton, S. W., Latreille, P., Porwollik, S., Sabo, A., Meyer, R., Bieri, T., Ozersky, P., McLellan, M., Harkins, C. R., Wang, C., Nguyen, C., Berghoff, A., Elliott, G., Kohlberg, S., Strong, C., Du, F., Carter, J., Wilson, R.K. (2004). Comparison of genome degradation in Paratyphi A and Typhi, human-restricted serovars of *Salmonella enterica* that cause typhoid. Nat Genet, 36(12), 1268–1274. https://doi.org/10.1038/ng1470

Moreau, K., Lacas-Gervais, S., Fujita, N., Sebbane, F., Yoshimori, T., Simonet, M., & Lafont, F. (2010). Autophagosomes can support *Yersinia pseudotuberculosis* replication in macrophages. Cell Microbiol, 12(8), 1108–1123. https://doi.org/10.1111/j.1462-5822.2010.01456.x

Niekamp, P., Scharte, F., Sokoya, T., Vittadello, L., Kim, Y., Deng, Y., Südhoff, E., Hilderink, A., Imlau, M., Clarke, C. J., Hensel, M., Burd, C. G., & Holthuis, J. C. M. (2022). Ca^2+^-activated sphingomyelin scrambling and turnover mediate ESCRT-independent lysosomal repair. Nat Commun, 13(1), 1875. https://doi.org/10.1038/s41467-022-29481-4

Noster, J., Chao, T. C., Sander, N., Schulte, M., Reuter, T., Hansmeier, N., & Hensel, M. (2019). Proteomics of intracellular *Salmonella enterica* reveals roles of *Salmonella* pathogenicity island 2 in metabolism and antioxidant defense. PLoS Pathog, 15(4), e1007741. https://doi.org/10.1371/journal.ppat.1007741

Ogawa, M., Yoshimori, T., Suzuki, T., Sagara, H., Mizushima, N., & Sasakawa, C. (2005). Escape of intracellular *Shigella* from autophagy. Science, 307(5710), 727–731. 15576571

Otten, E. G., Werner, E., Crespillo-Casado, A., Boyle, K. B., Dharamdasani, V., Pathe, C., Santhanam, B., & Randow, F. (2021). Ubiquitylation of lipopolysaccharide by RNF213 during bacterial infection. Nature, 594(7861), 111–116. https://doi.org/10.1038/s41586-021-03566-4

Paz, I., Sachse, M., Dupont, N., Mounier, J., Cederfur, C., Enninga, J., Leffler, H., Poirier, F., Prevost, M. C., Lafont, F., & Sansonetti, P. (2010). Galectin-3, a marker for vacuole lysis by invasive pathogens. Cell Microbiol, 12(4), 530–544. https://doi.org/10.1111/j.1462-5822.2009.01415.x

Reuter, T., Scharte, F., Franzkoch, R., Liss, V., & Hensel, M. (2021). Single cell analyses reveal distinct adaptation of typhoidal and non-typhoidal *Salmonella enterica* serovars to intracellular lifestyle. PLoS Pathog, 17(6), e1009319. https://doi.org/10.1371/journal.ppat.1009319

Röder, J., Felgner, P., & Hensel, M. (2021). Comprehensive Single Cell Analyses of the Nutritional Environment of Intracellular *Salmonella enterica*. Front Cell Infect Microbiol, 11, 624650. https://doi.org/10.3389/fcimb.2021.624650

Röder, J., & Hensel, M. (2020). Presence of SopE and mode of infection result in increased *Salmonella*-containing vacuole damage and cytosolic release during host cell infection by *Salmonella enterica*. Cell Microbiol, 22(5), e13155. https://doi.org/10.1111/cmi.13155

Schell, U., Simon, S., & Hilbi, H. (2016). Inflammasome Recognition and Regulation of the *Legionella* Flagellum. Curr Top Microbiol Immunol, 397, 161–181. https://doi.org/10.1007/978-3-319-41171-2_8

Schulte, M., Sterzenbach, T., Miskiewicz, K., Elpers, L., Hensel, M., & Hansmeier, N. (2019). A versatile remote control system for functional expression of bacterial virulence genes based on the *tetA* promoter. Int J Med Microbiol, 309(1), 54–65. https://doi.org/10.1016/j.ijmm.2018.11.001

Sepe, L. P., Hartl, K., Iftekhar, A., Berger, H., Kumar, N., Goosmann, C., Chopra, S., Schmidt, S. C., Gurumurthy, R. K., Meyer, T. F., & Boccellato, F. (2020). Genotoxic Effect of *Salmonella* Paratyphi A Infection on Human Primary Gallbladder Cells. mBio, 11(5). https://doi.org/10.1128/mBio.01911-20

Straley, S. C., & Harmon, P. A. (1984). *Yersinia pestis* grows within phagolysosomes in mouse peritoneal macrophages. Infect Immun, 45(3), 655–659. https://doi.org/10.1128/iai.45.3.655-659.1984

Striednig, B., Lanner, U., Niggli, S., Katic, A., Vormittag, S., Brulisauer, S., Hochstrasser, R., Kaech, A., Welin, A., Flieger, A., Ziegler, U., Schmidt, A., Hilbi, H., & Personnic, N. (2021). Quorum sensing governs a transmissive *Legionella* subpopulation at the pathogen vacuole periphery. EMBO Rep, 22(9), e52972. https://doi.org/10.15252/embr.202152972

Studer, P., Staubli, T., Wieser, N., Wolf, P., Schuppler, M., & Loessner, M. J. (2016). Proliferation of *Listeria monocytogenes* L-form cells by formation of internal and external vesicles. Nat Commun, 7, 13631. https://doi.org/10.1038/ncomms13631

Thurston, T. L., Wandel, M. P., von Muhlinen, N., Foeglein, A., & Randow, F. (2012). Galectin 8 targets damaged vesicles for autophagy to defend cells against bacterial invasion. Nature, 482(7385), 414–418. https://doi.org/10.1038/nature10744

Tinevez, J. Y., Perry, N., Schindelin, J., Hoopes, G. M., Reynolds, G. D., Laplantine, E., Bednarek, S. Y., Shorte, S. L., & Eliceiri, K. W. (2017). TrackMate: An open and extensible platform for single-particle tracking. Methods, 115, 80–90. https://doi.org/10.1016/j.ymeth.2016.09.016

TRC. (2015a, June, 3 2015). Lentivirus production of ShRNA, CRISPR, or ORF-pLX clones in 10 cm dishes or 6-well plates. Retrieved July, 4 2022 from https://portals.broadinstitute.org/gpp/public/resources/protocols

TRC. (2015b, June, 3 2015). Lentivirus production of ShRNA, CRISPR, or ORF-pLX clones in 10 cm dishes or 6-well plates. https://portals.broadinstitute.org/gpp/public/resources/protocols

Valenzuela, L. M., Hidalgo, A. A., Rodriguez, L., Urrutia, I. M., Ortega, A. P., Villagra, N. A., Paredes-Sabja, D., Calderon, I. L., Gil, F., Saavedra, C. P., Mora, G. C., & Fuentes, J. A. (2015). Pseudogenization of sopA and sopE2 is functionally linked and contributes to virulence of Salmonella enterica serovar Typhi. Infect Genet Evol, 33, 131–142. https://doi.org/10.1016/j.meegid.2015.04.021

Wadhwa, N., & Berg, H. C. (2022). Bacterial motility: machinery and mechanisms. Nat Rev Microbiol, 20(3), 161–173. https://doi.org/10.1038/s41579-021-00626-4

Yoshikawa, Y., Ogawa, M., Hain, T., Yoshida, M., Fukumatsu, M., Kim, M., Mimuro, H., Nakagawa, I., Yanagawa, T., Ishii, T., Kakizuka, A., Sztul, E., Chakraborty, T., & Sasakawa, C. (2009). *Listeria monocytogenes* ActA-mediated escape from autophagic recognition. Nat Cell Biol, 11(10), 1233–1240. https://doi.org/10.1038/ncb1967

